# P-Rex1 Limits the Agonist-Induced Internalisation of GPCRs Independently of its Rac-GEF Activity

**DOI:** 10.1101/2025.04.14.648762

**Authors:** Martin J. Baker, Elizabeth Hampson, Priota Islam, Ruben Pelaez Moral, Eve A. Maunders, Kirsti Hornigold, Elpida Tsonou, David C. Hornigold, Roderick E. Hubbard, Andrew J. Massey, Heidi C. E. Welch

## Abstract

The Rac-GEF P-Rex1 mediates GPCR signalling by activating the small GTPase Rac. We show here that P-Rex1 also controls GPCR trafficking. P-Rex1 inhibits the agonist-stimulated internalisation of the GPCR S1PR1 independently of its Rac-GEF activity, through its PDZ, DEP and IP4P domains. P-Rex1 also limits the agonist-induced trafficking of CXCR4, PAR4, and GLP1R, but does not control steady-state GPCR levels, nor the agonist-induced internalisation of the RTKs PDGFR and EGFR. P-Rex1 blocks the phosphorylation required for GPCR internalisation. P-Rex1 binds Grk2, both *in vitro* and in cells, but does not appear to regulate Grk2 activity. We propose that P-Rex1 limits the agonist-induced internalisation of GPCRs through its interaction with Grk2 to maintain high levels of active GPCR at the plasma membrane. Therefore, P-Rex1 plays a dual role in promoting GPCR responses, by controlling GPCR trafficking through an adaptor function as well as by mediating GPCR signalling through its Rac-GEF activity.

**Highlights:**

- P-Rex1 controls GPCR trafficking, independently of its Rac-GEF activity
- P-Rex1 limits the agonist-induced internalisation of S1PR1, CXCR4, PAR4 and GLP1R
- P-Rex1 does not control steady-state GPCR levels, or PDGFR and EGFR trafficking
- P-Rex1 binds Grk2 and inhibits the phosphorylation required for GPCR internalisation

**eTOC blurb:** P-Rex1 activates Rac downstream of GPCRs to regulate processes ranging from innate immunity to neuronal plasticity, its deregulation contributing to cancer. Here, Baker et al. show that P-Rex1 also controls GPCR trafficking, limiting agonist-induced GPCR internalisation through an adaptor function. Thus, P-Rex1 promotes GPCR responses in a dual manner.

**Graphical Abstract:** 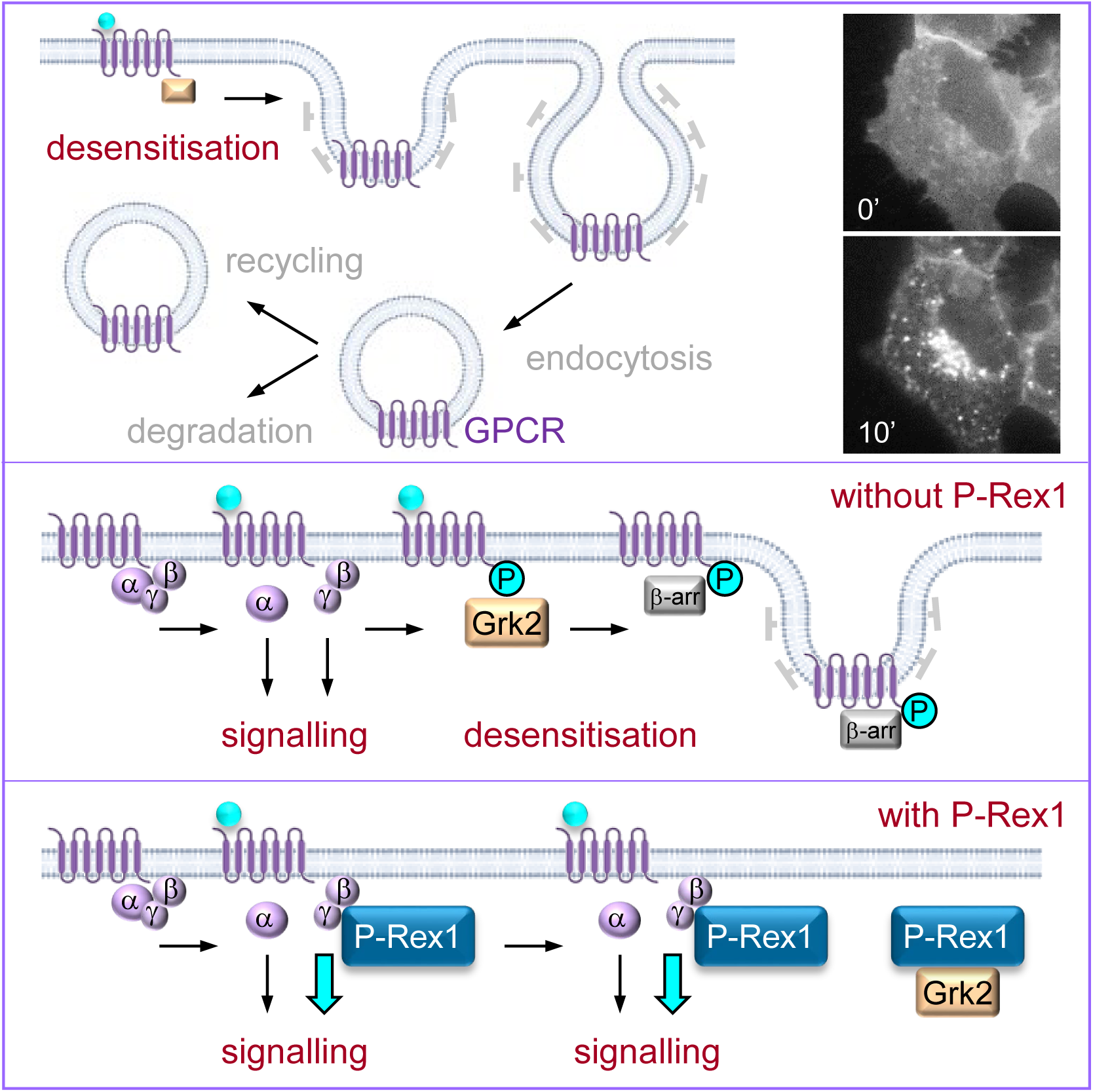

## Introduction

P-Rex1 is a guanine-nucleotide exchange factor (GEF) for Rac-type small GTPases which is widely expressed and plays important roles in the immune and nervous systems ^1−4^. In the immune system, P-Rex1 is required for a range of pro-inflammatory and immune functions, including the recruitment of leukocytes to sites of inflammation and infection, and for the clearance of bacteria ^3,5−8^. In the nervous system, P-Rex1 is required for synaptic plasticity, fine motor skills, and social recognition ^9−11^. Furthermore, P-Rex1 is required for melanoblast migration during development, controlling skin pigmentation ^12^. Deregulation of P-Rex1 levels contributes to a range of diseases. P-Rex1 overexpression is seen in many types of cancers, including breast cancer and melanoma, promoting tumour initiation, growth and/or invasiveness ^12−19^. Loss of P-Rex1 is associated with autism spectrum disorders ^11^. Furthermore, in mice, P-Rex1 promotes pulmonary fibrosis ^20^ and diet-induced non-alcoholic fatty liver disease ^21^.

All physiological and pathophysiological functions of P-Rex1 are either known or assumed to be mediated through its catalytic Rac-GEF activity. P-Rex1 activates all isoforms of Rac (Rac1, Rac2, Rac3, RhoG) upon itself being activated synergistically by the phosphoinositide 3-kinase (PI3K)-generated lipid second messenger phosphoinositide 3,4,5-trisphosphate (PIP_3_) and by the Gβγ subunits of heterotrimeric G proteins which are released upon activation of G protein-coupled receptors (GPCRs) ^1^. P-Rex1 has an N-terminal catalytic DH domain which serves to activate Rac, a PH domain, two DEP and two PDZ domains, and a C-terminal half similar to inositol polyphosphate 4-phosphatase (IP4P) but without phosphatase activity ^1,3^. Residues E56 and N238 in the DH domain are required for the interaction with Rac1 and for catalytic activity ^4,22^. PIP_3_ binds to the PH domain and Gβγ bind to the DH domain and to C-terminal domains, whereas the DEP, PDZ and IP4P domains serve to keep the protein in an autoinhibited confirmation prior to cell stimulation ^22−30^.

In addition to P-Rex1, the P-Rex family also comprises P-Rex2, a Rac-GEF with the same domain structure and regulated in the same manner by PIP_3_ and Gβγ ^9,31,32^. However, unlike P-Rex1, P-Rex2 has a known adaptor function, binding and inhibiting the tumour suppressor PTEN which converts PIP_3_ to PI(4,5)P_2_, and thus indirectly stimulating PI3K pathway activity, independently of its catalytic Rac-GEF activity ^33−35^. This adaptor function is specific to P-Rex2, as P-Rex1 cannot bind PTEN ^33^.

GPCRs are the largest family of cell surface receptors, characterised by their seven transmembrane spanning regions ^36^. They signal in response to a vast array of stimuli, ranging from photons to proteins, peptides and lipids. Ligand binding induces a conformational change, which activates the receptor-coupled heterotrimeric G protein, consisting of Gα and Gβγ ^37^. The Gα subunit is a GTPase, and the ligand-bound GPCR acts as its GEF, leading to GTP-loading of Gα and release of the Gβγ dimer. Gα-GTP and Gβγ then interact with their respective effector proteins to elicit downstream signalling ^37^.

GPCRs can be classified according to the type of Gα they couple to, Gα_s_ Gαi, Gα_q_ or Gα_12/13_. Very generally, Gα_s_ signals to increase adenylyl cyclase activity and thus cellular cAMP levels, Gαi inhibits adenylyl cyclase, Gα_q_ activates phospholipase-C-β (PLCβ) to stimulate Ca^2+^ signalling, and Gα_12/13_ activates Rho-GEFs to regulate cytoskeletal organisation ^38^. However, all Gα proteins signal through multiple pathways. Like Gα, Gβγ proteins also regulate a variety of signalling pathways, including the P-Rex Rac-GEF, PLCβ and PI3Kγ pathways ^39^.

The desensitisation of activated GPCRs occurs through the agonist-induced internalisation of the receptors ^40,41^. G protein-coupled receptor kinases (GRKs) recognise active GPCRs and phosphorylate serine or threonine residues in their cytoplasmic C-terminus. β-arrestin is recruited to the phosphorylated receptor, which sterically hinders the coupling between GPCR and heterotrimeric G protein and recruits clathrin adaptor AP2, Arf-GTPase Arf6 and clathrin, leading to clathrin-mediated endocytosis of the GPCR. This usually terminates the GPCR signal, although some internalised GPCRs may continue to signal intracellularly. The internalised GPCRs are then either recycled back to the cell membrane following dephosphorylation, or they are transported to lysosomes for degradation.

The GRK family, which comprises seven members, serve to phosphorylate active GPCRs for desensitisation ^42^. Grk2, the prototype of the family, is a ubiquitously expressed kinase which carries an N-terminal regulator of G proteins signalling homology (RH) domain, central catalytic kinase domain and C-terminal PH domain ^43,44^. Binding of Gβγ to the PH domain upon GPCR activation recruits Grk2 to the plasma membrane, enabling the catalytic domain to phosphorylate the GPCR ^42^.

Here, we investigated roles of P-Rex1 in GPCR trafficking, and show that P-Rex1 binds directly to GRK2 and limits the agonist-induced internalisation of GPCRs through an adaptor function.

## Results

### P-Rex1 limits the agonist-induced internalisation of the GPCR S1PR1 independently of its catalytic Rac-GEF activity

Sphingosine 1-phosphate receptor 1 (S1PR1) is a widely expressed GPCR for sphingosine 1-phosphate (S1P), with essential roles in vascular and neuronal development and pleiotropic other functions including immune cell migration ^45^. Upon S1P stimulation, S1PR1 signals through to Gα_i_ and is then internalised by clathrin-mediated endocytosis to switch off its signalling ^46^. To study S1PR1 trafficking, we generated HEK293 cells which stably express S1PR1-GFP (HEK293-S1PR1 cells), similar to a neuronal PC12-S1PR1 cell line we previously established ^47^. Widefield fluorescence microscopy showed that S1PR1-GFP is localised at the plasma membrane of serum-starved cells, as judged by the sheet-like staining typically seen in widefield imaging, and is internalised into vesicles upon S1P stimulation in a dose- and time dependent manner, with an EC_50_ of 5 nM S1P after 30 min and with 50% internalisation after 17 min in response to 10 nM S1P **(Supplemental Figure 1A-D)**. We assessed the localisation of S1PR1-GFP by two image analysis methods, a semi-quantitative method comparing images to a panel of standard images **(Supplemental Figure 1A, B, E)**, and a quantitative method using Volocity image analysis ^48^ **(Supplemental Figure 2A-C)**, which gave similar results.

To investigate whether P-Rex1 plays a role in S1PR1 trafficking, we expressed EE-tagged P-Rex1 transiently in HEK293-S1PR1 cells, stimulated the cells with 10 nM S1P, fixed, and imaged them for EE-P-Rex1 and for the localisation of S1PR1-GFP. EE-P-Rex1 expression limited the S1P-stimulated internalisation of the receptor **(Figure 1A)**. The same results were obtained by live-cell imaging. HEK293-S1PR1 cells showed the expected robust internalisation of S1PR1-GFP from the plasma membrane into intracellular vesicles in response to S1P **(Supplemental Movie 1)**, whereas mCherry-P-Rex1 reduced this receptor internalisation **(Supplemental Movie 2)**. Therefore, P-Rex1 limits the agonist-induced internalisation of the GPCR S1PR1. In contrast, S1PR1 localisation was normal without S1P stimulation **(Figure 1A and Supplemental Figure 2B)**, suggesting that P-Rex1 inhibits agonist-dependent but not steady-state receptor trafficking. Furthermore, P-Rex1 did not affect the total expression level of the GPCR **(Supplemental Figure 2D)**.

**Figure 1.**
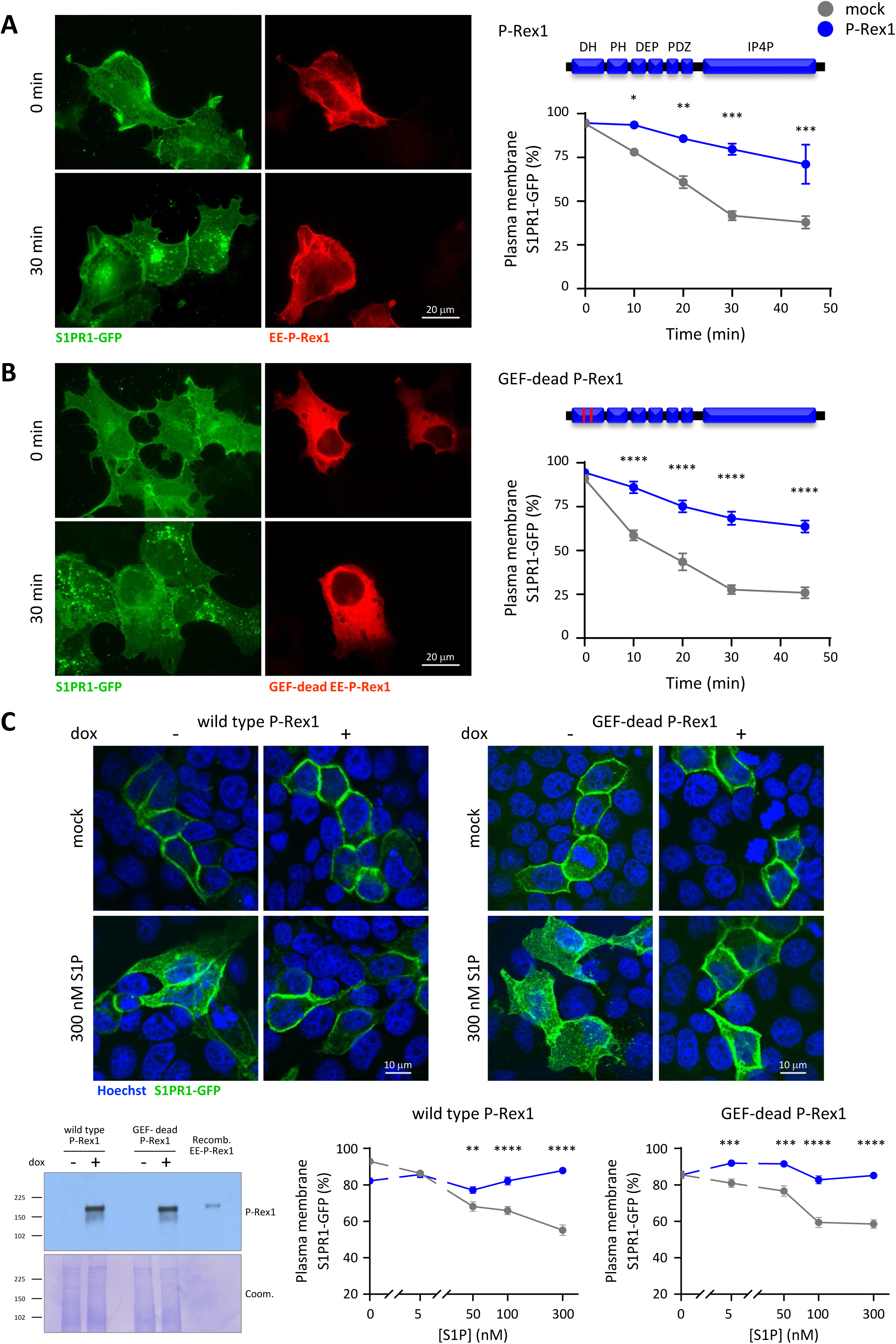
P-Rex1 limits the S1P-dependent internalisation of S1PR1 independently of its catalytic Rac-GEF activity. **(A, B)** Wild type and GEF-dead P-Rex1 limit the S1P-induced internalisation of S1PR1. HEK293-S1PR1 cells, which express S1PR1-GFP, were transfected with wild type (A) or GEF-dead (B) EE-P-Rex1 (blue symbols), or mock transfected (grey symbols), serum-starved, stimulated with 10 nM S1P for the indicated periods of time, fixed, and stained with EE antibody. Representative images show cells stimulated for 0 and 30 min, respectively. S1PR1-GFP localisation at the plasma membrane was quantified by comparison to standard images (see Supplemental Figure 1). Alternative quantification of the same data by Volocity image analysis is shown in Supplemental Figure 2B-C. Data are mean ± SEM of 3 independent experiments. Statistics are two-way ANOVA with Sidak’s multiple comparisons correction; stars denote differences between genotypes for each time point. **(C)** Inducible expression of wild type or GEF-dead P-Rex1 inhibits the S1P-stimulated internalisation of S1PR1. MDCK cells with doxycycline (dox)-inducible expression of wild type or GEF-dead P-Rex1 were treated with (blue) or without (grey) 1 µg/ml dox for 24 h, serum-starved, and stimulated with the indicated concentrations of S1P for 10 min, fixed, stained with Hoechst 33342, and imaged by confocal fluorescence microscopy. Representative confocal images are shown. Quantification was done as in (A, B). Data are mean ± SEM of 66-95 cells per condition. Statistics are two-way ANOVA with Sidak’s multiple comparisons correction; stars denote differences between genotypes for each S1P concentration. The western blot shows the dox-induced expression of wild type or GEF-dead P-Rex1 under the same conditions. Recombinant EE-P-Rex1 was loaded as a control. Coomassie staining was used as a control for protein loading.

To determine if suppression of GPCR trafficking depends on the catalytic Rac-GEF activity of P-Rex1, we expressed GEF-dead P-Rex1, which contains point mutations E56A and N238A in the DH domain which abolish GEF-activity ^22,25^. As previously observed for wild type P-Rex1, S1PR1-GFP was retained at the plasma membrane of GEF-dead EE-P-Rex1 expressing cells stimulated with S1P **(Figure 1B and Supplemental Figure 2C)**. Hence, P-Rex1-dependent control of GPCR trafficking is independent of its catalytic Rac-GEF activity.

To test a different cell system, we used MDCK cells with doxycycline (dox)-inducible expression of wild type or GEF-dead P-Rex1 ^22,49^ **(Figure 1C)**, together with transient expression of S1PR1-GFP, and performed confocal fluorescence microscopy. Without dox-induction, S1PR1-GFP was seen as a ring at the cell periphery, as typically seen for plasma membrane-localised proteins by confocal microscopy, and the receptor was internalised into intracellular vesicles upon S1P stimulation in a dose-dependent manner as expected. Expression of wild type or GEF-dead P-Rex1 inhibited this receptor internalisation, as previously seen in HEK293-S1PR1 cells **(Figure 1C)**.

Therefore, P-Rex1 inhibits the agonist-induced internalisation of S1PR1-GFP independently of its catalytic Rac-GEF activity. This is the first Rac-GEF activity independent function of P-Rex1.

### P-Rex1 deficiency increases the agonist-induced internalisation of S1PR1

To investigate whether endogenous P-Rex1 plays a role in the S1P-stimulated internalisation of S1PR1-GFP, we used wild type (*Prex1^+/+^*) and two clones of P-Rex1-deficient (*Prex1^−/−^*) PC12-S1PR1 cells ^47^. The cells were serum-starved, stimulated with S1P, and the localisation of PC12-S1PR1 was assessed by confocal fluorescence microscopy **(Figure 2A-C and Supplemental Figure 1E)**. P-Rex1 deficiency increased S1P-induced receptor internalisation, whereas the steady state-cell surface level of the receptor was normal **(Figure 2B, C)**. Hence, P-Rex1 expression limits and P-Rex1 deficiency promotes the agonist-induced internalisation of S1PR1-GFP.

**Figure 2.**
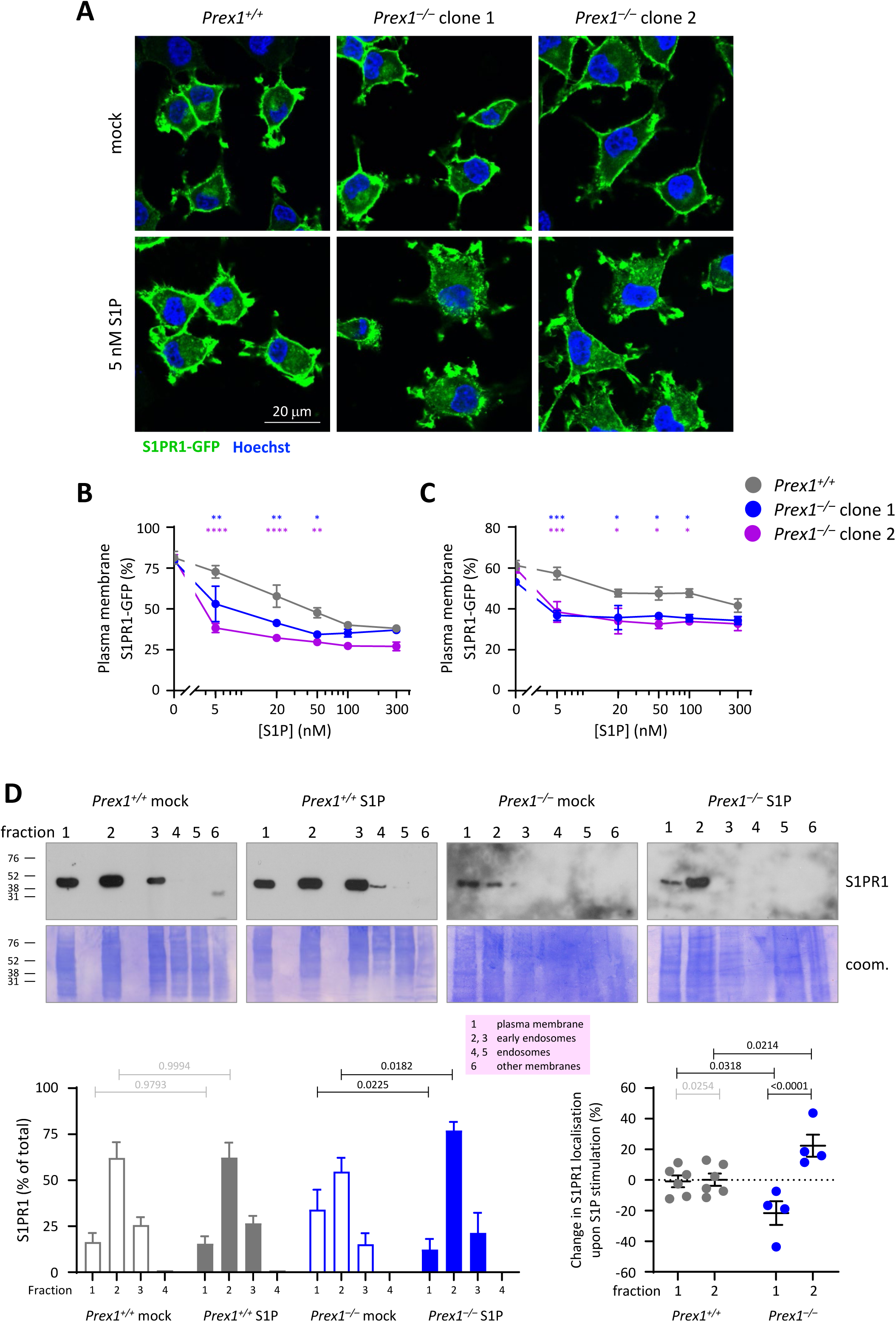
P-Rex1 deficiency promotes the S1P-dependent internalisation of S1PR1. (A-C) Wild type PC12-S1PR1 (*Prex1^+/+^*, grey symbols) and Prex1-deficient PC12-S1PR1 cells clone 1 (blue) and clone 2 (purple) were serum-starved, stimulated with the indicated concentrations of S1P for 10 min, fixed, stained with Hoechst 33342, and imaged by confocal microscopy. (A) Representative confocal images. S1PR1 localisation at the plasma membrane was quantified by (B) comparison to standard images (see Supplemental Figure 1E) or by (C) CellProfiler analysis (see Supplemental Figure 2A). Data in (B) and (C) are mean ± SEM of ≥3 independent experiments; the same experiments were analysed by both methods. Statistics are two-way ANOVA, with Sidak’s multiple comparisons correction; stars denote differences between genotypes for each S1P concentration. **(D)** Fractionation of PC12-S1PR1 cells. *Prex1^+/+^* (grey) and *Prex1^−/−^* (blue) PC12-S1PR1 cells were serum-starved and stimulated with 5 nM S1P, or mock-stimulated, for 10 min. Detergent-free cell lysates were fractionated by discontinuous OptiPrep density gradient (see Supplemental Figure 3), and proteins from each fraction analysed by western blotting with S1PR1 antibody. All of fractions 1-5 and 50% of fraction 6 were loaded. Representative western blots are shown. Coomassie staining was used to show total protein. Lower left: The amount of endogenous S1PR1 per fraction was quantified by Fiji densitometry. Lower right: change in S1PR1 localisation in fractions 1 and 2 upon S1P stimulation. Data are mean ± SEM from 4-6 independent experiments. Statistics are two-way ANOVA with Sidak’s multiple comparisons correction; black p-values denote significant differences, grey p-values are not significant.

To assess whether P-Rex1 controls the trafficking of endogenous S1PR1, and for an alternative method of quantifying receptor internalisation, we adapted a previously described cell fractionation method to separate endosomes from the plasma membrane in PC12 cells ^50^, using ultracentrifugation of detergent-free PC12-S1PR1 cell lysates on an OptiPrep density-gradient followed by western blotting. This provided a K-Ras-enriched plasma-membrane fraction (fraction 1), EEA1-enriched early endosome fractions (2, 3), and a Rab5-enriched endosome fraction (5) **(Supplemental Figure 3A-C)**. Fractionation of serum-starved *Prex1^+/+^* and *Prex1^−/−^* PC12-S1PR1 cells showed that endogenous S1PR1 was mainly localised in the plasma membrane and early endosome fractions of both genotypes. Upon stimulation with a low concentration of S1P (5 nM), 22% of the total cellular S1PR1 translocated from the plasma membrane into the early endosome fraction in *Prex1^−/−^* PC12-S1PR1 cells, whereas receptor localisation remained unchanged in *Prex1^+/+^* PC12-S1PR1 cells **(Figure 2D)**. This extent of internalisation of endogenous S1PR1 measured by fractionation was identical to that of S1PR1-GFP assessed by imaging at that concentration of S1P. To test the specificity of S1PR1 internalisation, we also determined the localisation of the receptor tyrosine kinase (RTK) EGFR. Endogenous EGFR was localised throughout plasma membrane and endosomal fractions in both *Prex1^+/+^* and *Prex1^−/−^*PC12-S1PR1 cells, and S1P stimulation did not affect its localisation **(Supplemental Figure 3D)**. Therefore, P-Rex1 limits the agonist-induced internalisation of endogenous S1PR1.

### Large portions of the P-Rex1 protein are required for its control of GPCR trafficking

To identify which P-Rex1 domains are required for the control of GCPR trafficking, we expressed various P-Rex1 mutants ^22,51^. EE-P-Rex1 ΔPH inhibited the S1P-induced internalisation of S1PR1-GFP similar to wild type P-Rex1 **(Figure 3A and Supplemental Figure 4A)**, suggesting that the PH domain is dispensable. In contrast, EE-P-Rex1 ΔPDZ, ΔDEP, and ΔIP4P mutants had no effect on the trafficking of S1PR1-GFP **(Figure 3B-D and Supplemental Figure 4B-D)**. Therefore, large portions of the P-Rex1 protein, including the DEP, PDZ and IP4P domains, are required for its ability to inhibit GPCR trafficking. Furthermore, it was previously suggested that the isolated PDZ domains of P-Rex1 can interact with S1PR1 ^52^. To investigate, we expressed myc-P-Rex1 iPDZ in HEK293-S1PR1 cells. This had no effect on the S1P-stimulated trafficking of S1PR1-GFP **(Supplemental Figure 5A)**, suggesting that isolated PDZ domain tandem is not sufficient for the control of S1PR1 trafficking.

**Figure 3.**
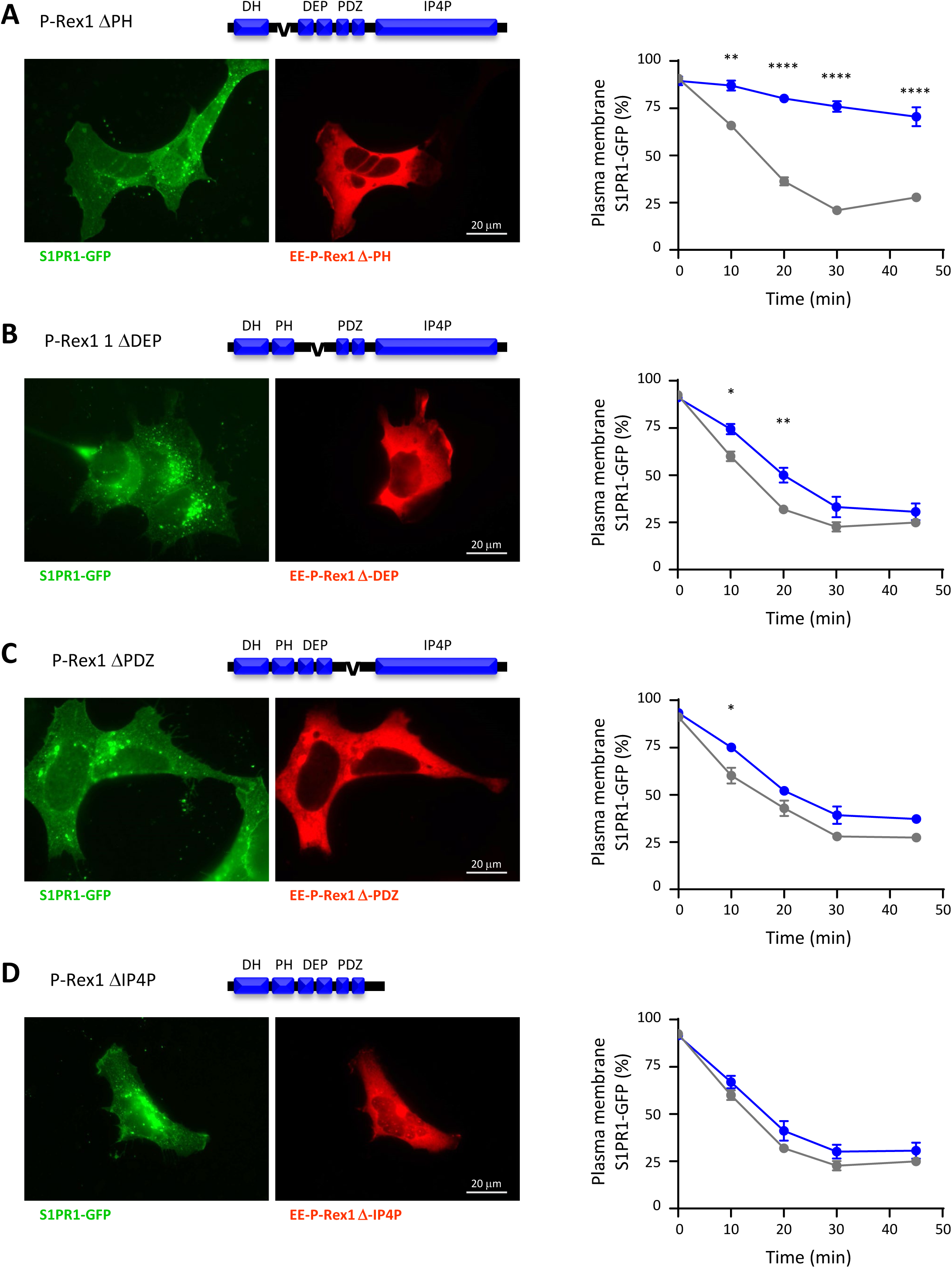
The DEP, PDZ and IP4P domains of P-Rex1 are required for the inhibition of S1PR1 internalisation. (A-D) HEK293-S1PR1 cells were transfected with (A) EE-P P-Rex1 ΔPH, (B) EE-P P-Rex1 ΔDEP, (C) EE-P P-Rex1 ΔPDZ, or EE-P-Rex1 ΔIP4P (D) (blue symbols), or mock-transfected (grey symbols), serum-starved, stimulated with 10 nM S1P for the indicated periods of time, fixed and stained for EE. Representative images show cells stimulated with S1P for 30 min. S1PR1-GFP localisation at the plasma membrane was quantified by comparison to standard images (see Supplemental Figure 1). Alternative quantification of the same data by Volocity image analysis is shown in Supplemental Figure 4. Data are mean ± SEM of 3 independent experiments for each mutant; statistics are two-way ANOVA with Sidak’s multiple comparisons correction; stars denote differences between cells with and without P-Rex1 mutant for each time point.

To investigate whether the P-Rex1 homologue P-Rex2 plays a similar role in GPCR trafficking to P-Rex1, we expressed myc-P-Rex2 in HEK293-S1PR1 cells. P-Rex2 had a similar effect on S1P-induced S1PR1-GFP trafficking as P-Rex1 **(Supplemental Figure 5B)**. Therefore, both P-Rex family members limit agonist-induced GPCR internalisation.

### P-Rex1 limits the agonist-induced internalisation of a range of GPCRs but not that of RTKs

To investigate if other GPCRs are affected by P-Rex1, in addition to S1PR1, we selected candidate GPCRs to cover a range of classes that couple to different types of heterotrimeric G protein and expressed them in MDCK cells with dox-inducible expression of wild type or GEF-dead P-Rex1. The GPCRs included CXCR4 which, like S1PR1, is Gαi-coupled ^53^, Gα_s_-coupled Glucagon-Like Peptide-1 Receptor (GLP1R) ^54^, and Gα_q/12/13_-coupled Protease-Activated Receptor 4 (PAR4) ^55^. Each GPCR was assessed by confocal fluorescence microscopy, using stimulation with its own agonist, after pilot experiments to determine appropriate agonist concentrations and timing.

MDCK cells expressing CXCR4-LSSmOrange were treated with dox to induce P-Rex1 expression, or mock-treated, serum-starved, and stimulated with 25 nM SDF1α for various periods of time. The localisation of CXCR4-LSSmOrange was more cytoplasmic than previously observed for S1PR1, and even partially nuclear, which has been observed before ^56^, but SDF1α stimulation induced the expected dose-dependent internalisation of CXCR4-LSSmOrange from the plasma membrane into intracellular vesicles. Both wild type and GEF-dead P-Rex1 inhibited this SDF1α-stimulated internalisation of CXCR4-LSSmOrange **(Figure 4A)**. Similarly, MDCK cells expressing GLP1R-mCherry were stimulated for 10 min with various concentrations of Glucagon-Like Peptide-1 (GLP-1) ^57^. Again, the localisation of GLP1R-mCherry was more cytoplasmic than previously observed for S1PR1, but GLP-1 stimulation caused the expected dose-dependent internalisation of the receptor, and both wild type and GEF-dead P-Rex1 inhibited this GLP-1 stimulated internalisation **(Figure 4B)**. MDCK cells expressing PAR4-mCherry were stimulated with 500 µM AY-NH2 ^58^ for various periods of time. AY-NH2 stimulation caused the expected internalisation of PAR4-mCherry, and again, both wild type and GEF-Dead P-Rex1 inhibited this internalisation **(Figure 4C)**. Hence, P-Rex1 inhibits the agonist-induced internalisation of all GPCRs tested, regardless of the type of heterotrimeric G protein the GPCRs couple to, and in a GEF-activity independent manner.

**Figure 4.**
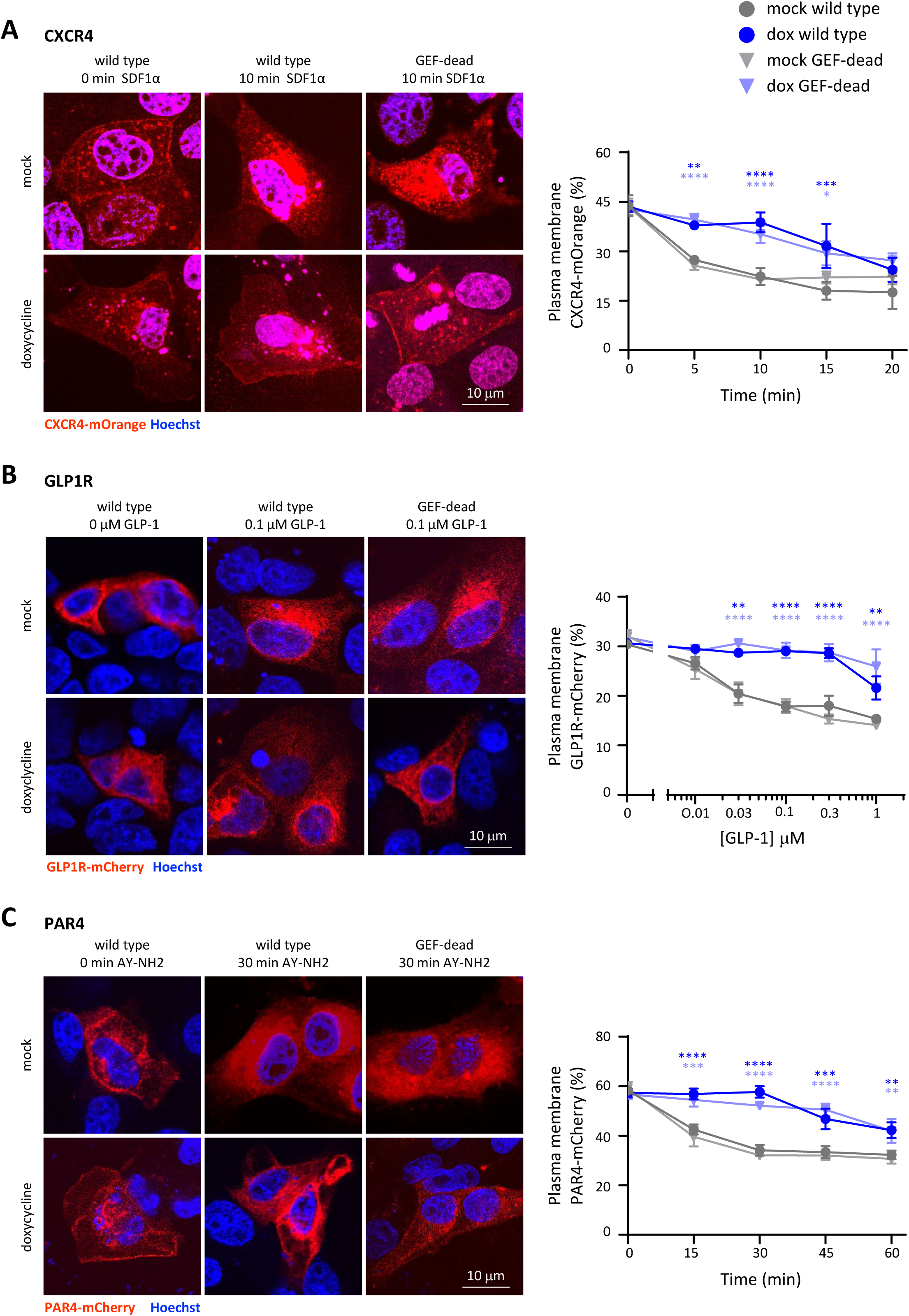
Expression of P-Rex1 inhibits the agonist-induced internalisation of CXCR4, GLP1R and PAR1, independently of its catalytic Rac-GEF activity. **(A)** CXCR4. CXCR4-LSSmOrange was expressed in MDCK cells with dox-inducible wild type (circles) or GEF-dead (triangles) P-Rex1. Cells were treated with 1 µg/ml dox (blue symbols) for 24 h, or mock-treated (grey symbols), serum starved, stimulated with 25 nM SDF1α for the indicated periods of time, fixed, stained with Hoechst 33342, and imaged by confocal fluorescence microscopy. Representative confocal images are shown. CXCR4-LSSmOrange localisation at the plasma membrane was quantified by comparison to standard images (see Supplemental Figure 1). **(B)** GLP1R. MDCK cells were treated as in (A) except that GLP1R-mCherry was expressed and cells were stimulated with the indicated concentrations of GLP-1 for 10 min. GLP1R-mCherry localisation was quantified as in (A). **(C)** PAR4. MDCK cells were treated as in (A, B) except that PAR4-mCherry was expressed and cells were stimulated with 500 µM AY-NH2 for the indicated periods of time. PAR4-mCherry localisation was quantified as in (A). Data in (A-C) are mean ± SEM of three independent experiments for each receptor. Statistics are two-way ANOVA with Sidak’s multiple comparisons correction; stars denote differences between mock and dox conditions.

To determine if P-Rex1 also controls the trafficking of other classes of receptor, we tested the agonist-induced internalisation of the RTKs EGFR and PDGFRβ. MDCK cells expressing EGFR-GFP were serum-starved and stimulated with 100 ng/ml EGF ^59^ for various periods of time. EGFR-GFP was largely localised at the plasma membrane of serum-starved cells, and EGF stimulation caused the expected internalisation of the receptor, but the agonist-induced internalisation of EGFR was not affected by wild type or GEF-dead P-Rex1 **(Supplemental Figure 6A)**. Similarly, PDGFRβ-GFP was also largely localised at the plasma membrane, and stimulation with 50 ng/ml PDGF ^60^ induced the internalisation of the receptor. Again, this internalisation was not affected by the expression of wild type or GEF-dead P-Rex1 **(Supplemental Figure 6B)**. Hence, P-Rex1 controls the trafficking of a range of GPCRs, in a GEF-activity independent manner, but does not control the trafficking of RTKs. Of note, we previously measured the cell surface levels of the L-selectin and of the β2 integrins LFA1 and Mac1 in *Prex1^−/−^* mouse neutrophils, which were also normal ^7,61^. Hence, P-Rex1 regulates the trafficking of GPCRs but not of a range of other receptor classes.

### P-Rex1 inhibits the phosphorylation which is required for GPCR internalisation

The first step in the agonist-induced internalisation of GPCRs is the phosphorylation of the GPCR at its C-terminal tail, mainly by GRKs ^42^. To investigate the effects of P-Rex1 on the phosphorylation of S1PR1, we used mass spectrometry. HEK293-S1PR1 cells expressing EE-P-Rex1 were serum-starved and stimulated with 10 nM S1P, or mock-stimulated, and S1PR1-GFP was immunoprecipitated from total lysates. Phosphorylated and non-phosphorylated peptides of the C-terminal tail, in particular a peptide encompassing S351, a residue critical for S1PR1 internalisation, were identified via LC-MS. In control cells, S1P stimulation increased the phosphorylation of S1PR1-GFP on S351. This S1P-induced phosphorylation was inhibited by the expression of P-Rex1 **(Figure 5A)**. Hence, P-Rex1 inhibits the S1P-dependent phosphorylation of S1PR1 on S351, the first step in the process of clathrin-mediated endocytosis of the receptor.

**Figure 5.**
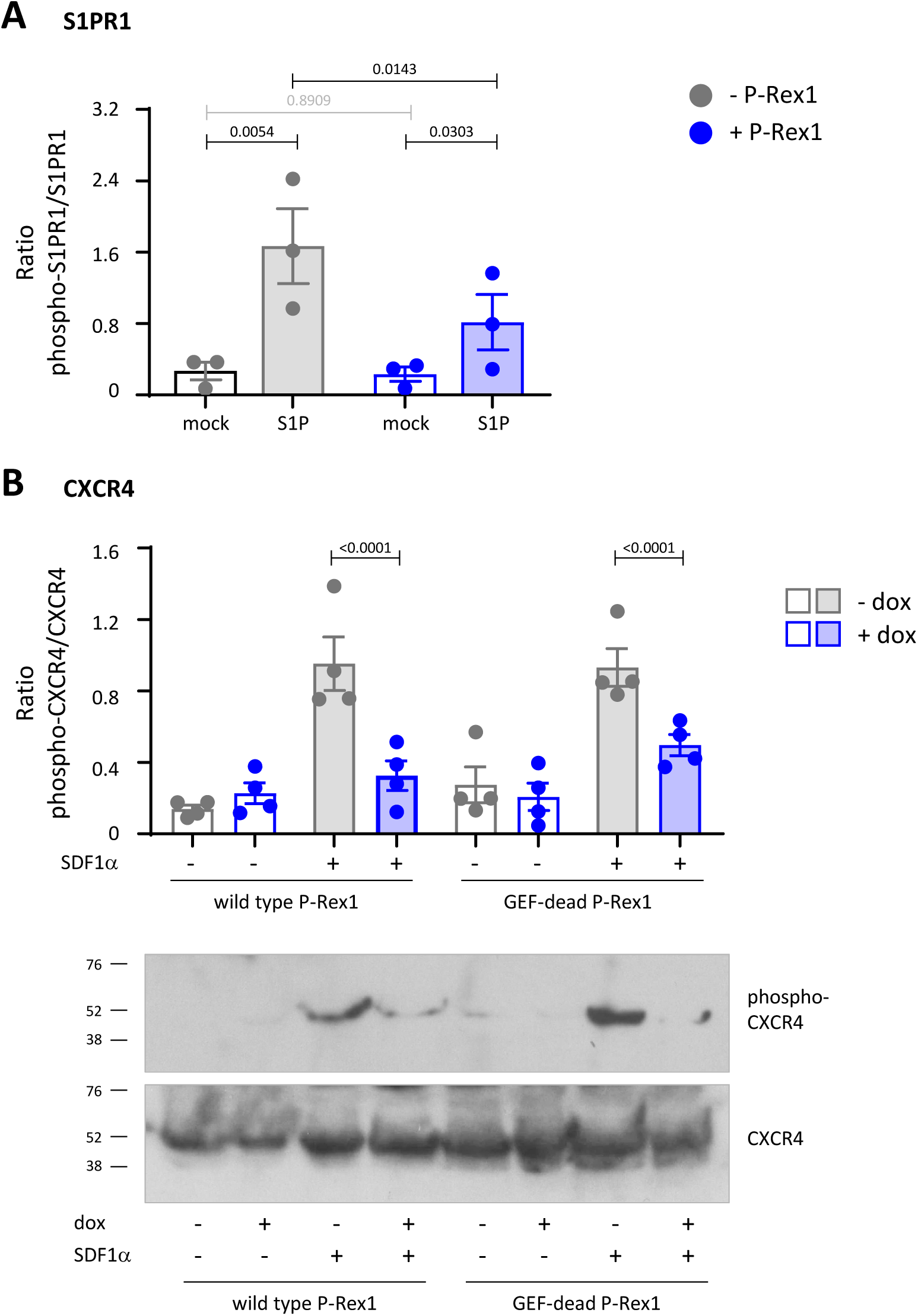
P-Rex1 inhibits the phosphorylation required for internalisation of S1PR1 and CXCR4. **(A)** P-Rex1 inhibits the S1P-stimulated phosphorylation of S1PR1-GFP at S351. S1PR1-GFP was immunoprecipitated from lysates of HEK293-S1PR1 cells that had been transfected with EE-P-Rex1 (blue symbols), or mock transfected (grey symbols), serum starved, and stimulated with 10 nM S1P for 10 min (filled bars), or mock-stimulated (open bars). Samples were analysed by LC-MS, targeted to phosphopeptides in the S1PR1 C-terminus. Data show the ratio of phosphorylated/non-phosphorylated S351 and are mean ± SEM of 3 independent experiments. Statistics are two-way ANOVA with Sidak’s multiple comparisons correction; black p-values denote significant differences, grey p-values are not significant. **(B)** Wild type and GEF-dead P-Rex1 inhibit the phosphorylation of CXCR4 at S3254/S325. MDCK cells were treated with 1 µg/ml dox for 24 h to induce the expression of wild type or GEF-dead P-Rex1 (blue), or were mock-treated (grey), serum-starved, and stimulated with 25 nM SDF1α for 10 min (filled symbols), or mock-stimulated (open symbols). Total cell lysates were western blotted with phospho-S324/S325-CXCR4 and total CXCR4 antibodies. Representative blots are shown. Blots were quantified by Fiji densitometry. Data show the ratio of phosphorylated/non-phosphorylated CXCR4 and are mean ± SEM of 4 independent experiments; statistics two-way ANOVA with Sidak’s multiple comparisons correction.

To investigate a different GPCR and test if limiting GPCR phosphorylation is a GEF-activity dependent role of P-Rex1, we assess the phosphorylation of endogenous CXCR4 in dox-inducible MDCK cells. The agonist-induced phosphorylation of residues S324 and S325 is required for the internalisation of CXCR4 ^62^. MDCK cells were induced with dox, or mock-treated, serum-starved, stimulated with 25 nM SDF1α for 10 min, or mock-stimulated, and total lysates western blotted for phospho-S324/S325 and total CXCR4. In control cells, SDF1α stimulation increased the phosphorylation of S324/S325, whereas the expression of wild type or GEF-dead P-Rex1 reduced this phosphorylation **(Figure 5B)**. Hence, P-Rex1 blocks the phosphorylation required for the agonist-induced internalisation of CXCR4, in a GEF-activity independent manner.

Gβγ proteins, which are released from Gα upon activation of GPCRs, can bind and activate P-Rex1. To investigate if Gβγ binding is involved in the effects of P-Rex1 on GPCR trafficking, we used gallein, a small molecule inhibitor of Gβγ interactions with effectors ^63^. We saw no effects of gallein on the S1P-dependent internalisation of S1PR1-GFP in pilot experiments, neither in the presence nor absence of P-Rex1 (data not shown), suggesting that the interaction of P-Rex1 with Gβγ plays no role in its ability to regulate GPCR trafficking.

To test if P-Rex1 interacts with GPCRs, HEK293-S1PR1 cells expressing myc-P-Rex1, or mock-transfected, were serum-starved, stimulated with 100 nM S1P, or mock-stimulated, and total lysates were subjected to immunoprecipitation with GFP or myc antibodies using various conditions of stringency. No interaction between P-Rex1 and S1PR1-GFP could be detected by western blotting of the IP samples under any of the conditions tested **(Supplemental Figure 7A,** and data not shown).

### P-Rex1 interacts with Grk2 in cells and *in vitro*

To investigate if P-Rex1 interacts with Grks, which carry out the C-terminal phosphorylation of GPCRs upon agonist-stimulation, we selected Grk2, the best-understood and arguably most important of these kinases. To test if P-Rex1 interacts with Grk2 *in vivo*, HEK293-S1PR1 cells expressing myc-P-Rex1 and/or flag-Grk2 were serum-starved, and Grk2 isolated from total lysates by immunoprecipitation with flag antibody. Myc-P-Rex1 co-immunoprecipitated with flag-Grk2 under these conditions **(Figure 6A)**, which shows that P-Rex1 constitutively associates with Grk2 in cells.

**Figure 6.**
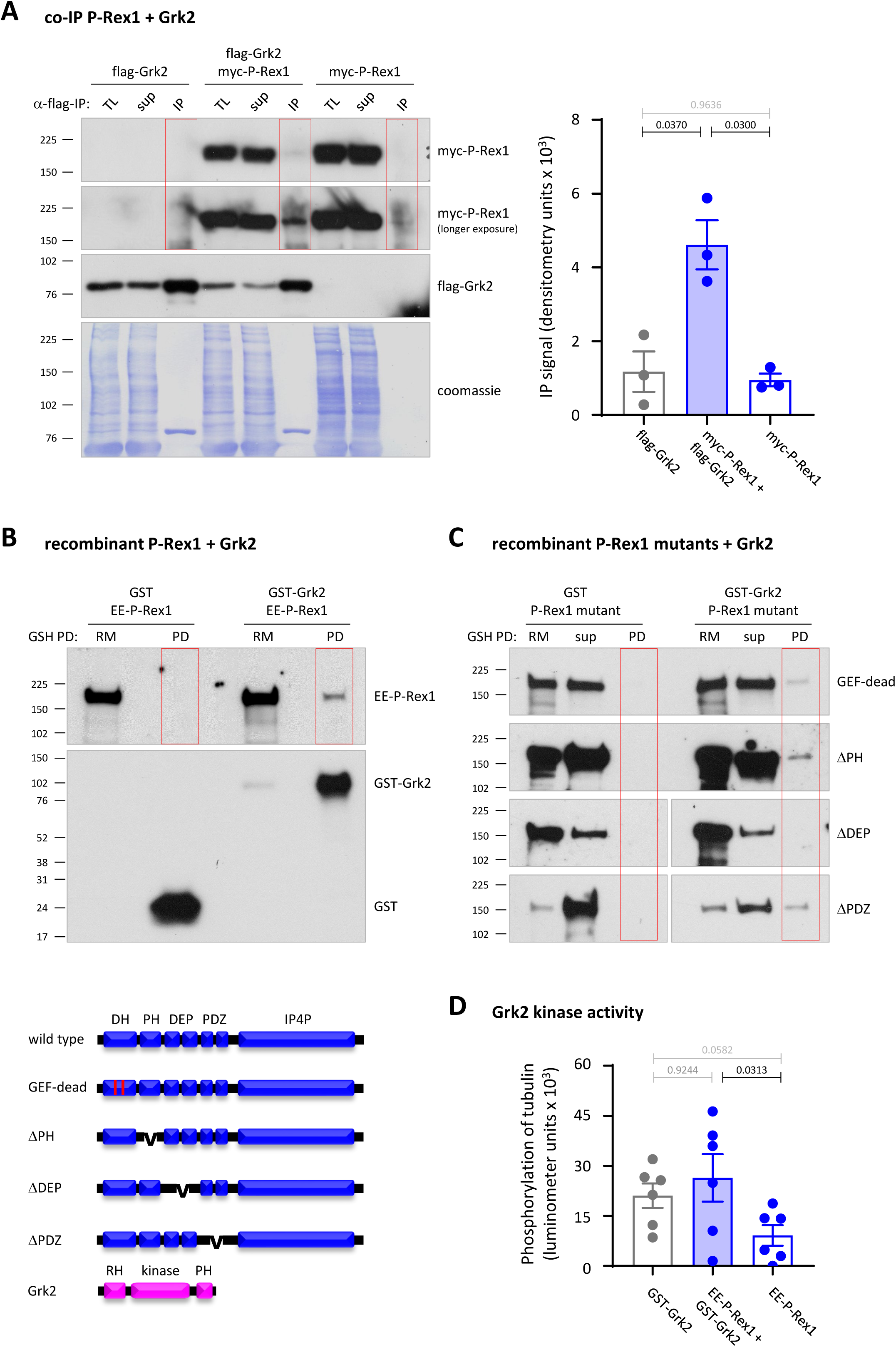
P-Rex1 interacts with Grk2. **(A)** P-Rex1 interacts with Grk2 *in vivo*. HEK293-S1PR1 cells expressing myc-P-Rex1 and/or flag-Grk2, were serum-starved, and total lysates were subjected to immunoprecipitation (IP) with flag antibody and analysed by western blotting with myc and flag antibodies. 1.5% of the total lysate (TL) and IP supernatant (sup) were loaded alongside all the IP sample (red boxes). Coomassie staining was used as a loading control. Representative western blots are shown. Blots were quantified by Fiji densitometry. Data are mean ± SEM of 3 independent experiments. Statistics are one-way ANOVA with Tukey’s multiple comparisons correction; black p-values denote significant differences, grey p-values are not significant. **(B, C)** P-Rex1 binds directly to Grk2 *in vitro* through its DEP domains. Wild type (B) or mutant (C) P-Rex1 proteins were incubated with GST or GST-Grk2, isolated using GSH-beads, and western blotted with P-Rex1 and GST antibodies. 10% of the reaction mix (RM) and, where indicated, pull down supernatant (sup) controls and all of the pull down (PD) sample were loaded. Blots are representative of 3 (B) and 2-3 (C) independent experiments per P-Rex protein. The quantification for (B) shown in Supplemental Figure 7. **(D**) P-Rex1 does not affect the kinase activity of Grk2. The kinase activity of GST-Grk2 was measured *in vitro*, with tubulin as the substrate, in the presence and absence of EE-P-Rex1. Data are mean ± SEM of 6 independent experiments. Statistics are one-way ANOVA with Tukey’s multiple comparisons correction; black p-values denote significant differences, grey p-values are not significant.

To test if P-Rex1 binds directly to Grk2 *in vitro*, and to determine if the interaction requires the GEF activity of P-Rex1, we incubated purified recombinant wild type and GEF-dead EE-P-Rex1 proteins with purified recombinant GST or GST-Grk2, isolated the GST-containing proteins by pull down with GSH beads, and analysed P-Rex1 binding by western blotting. EE-P-Rex1 bound to GST-Grk2 but not to GST **(Figure 6B and Supplemental Figure 7B)**. Hence, the constitutive binding between P-Rex1 and Grk2 is direct. The GEF activity was not required for this direct binding, as GEF-dead P-Rex1 also bound to flag-Grk2 **(Figure 6C)**. To test which domains of P-Rex1 are required for binding, we used purified recombinant P-Rex1 deletion mutants ^22,51^. The PH and PDZ domains were dispensable for Grk2 binding, but the DEP domain tandem was required **(Figure 6C)**. Hence, P-Rex1 binds Grk2 directly *in vitro* through its DEP domains, independently of its GEF activity.

We showed in Supplemental Figure 5B that P-Rex2 can regulate GPCR trafficking similar to P-Rex1. To test if P-Rex2 also binds Grk2, we generated human recombinant wild type and GEF-dead His-P-Rex2 proteins, the latter by introducing alanine mutations at Glu30 and Asn212 in the catalytic DH domain of P-Rex2, the equivalent residues to those in GEF-dead P-Rex1 ^22,51^, and purified the proteins from Sf9 cells **(Supplemental Figure 7C)**. For quality control, we used an *in vitro* Rac-GEF activity assay in the presence of the signalling lipid PIP3 to stimulate P-Rex2 activity, which confirmed that the purified wild type His-P-Rex2 is an active Rac-GEF and that P-Rex2^E30A,N212A^ is indeed GEF-dead **(Supplemental Figure 7C)**. We tested the binding of these proteins to GST-Grk2 in the same way as for P-Rex1. Both wild type and GEF-dead P-Rex2 bound to GST-Grk2 but not to GST **(Supplemental Figure 7D)**. Hence, like P-Rex1, P-Rex2 binds Grk2 directly *in vitro*, independently of its GEF activity.

Finally, to test if P-Rex1 affects Grk2 activity, we measured the kinase activity of GST-Grk2 *in vitro* with tubulin as the substrate, in the presence and absence of EE-P-Rex1. GST-Grk2 was able to phosphorylate tubulin, but EE-P-Rex1 had no effect on this kinase activity **(Figure 6D)**. Thus, while P-Rex1 constitutively interacts with Grk2, it does not seem to control Grk2 kinase activity.

Together our data show that P-Rex1 limits the agonist-induced internalisation of GPCRs, but not other types of receptors, interacts constitutively with the kinase Grk2 which is required for GPCR internalisation, and inhibits the C-terminal phosphorylation of GPCRs which is carried out by Grk2, all independently of its catalytic Rac-GEF activity and without obviously affecting the kinase activity of Grk2. We propose that P-Rex1 inhibits GPCR trafficking likely by physically hindering the access of Grk2 to the GPCR.

## Discussion

Our study shows that P-Rex1 inhibits the agonist-stimulated internalisation of GPCRs that switches-off GPCR signalling. This affects all GPCRs we tested, irrespective of the type of heterotrimeric G protein the receptors couple to. P-Rex1 inhibits the first step of the GPCR trafficking process, Grk-mediated receptor phosphorylation, which implies that all subsequent steps are also abrogated, as β-arrestin cannot be recruited to non-phosphorylated GPCRs, and the rest of the clathrin-mediated endocytosis machinery depends on β-arrestin recruitment ^40,41^. P-Rex1 did not affect the steady-state cell surface levels of GPCRs, nor total cellular GPCR levels, suggesting it plays no role in the constitutive trafficking or degradation of GPCRs. P-Rex1 is also unlikely to affect GPCR recycling back to the plasma membrane, as pilot experiments revealed normal localisation of the recycling endosome marker Rab11 in HEK293-S1PR1 cells (not shown). Furthermore, the localisation of EGFR was unaffected in S1P-stimulated *Prex1^−/−^* PC12-S1PR1 cells, and the agonist-stimulated internalisation of EGFR and PDGFR was normal. Therefore, P-Rex1 specifically limits the agonist-induced internalisation of GPCRs.

P-Rex1 controls GPCR trafficking independently of its Rac-GEF activity. This is the first description of a GEF-activity independent role of P-Rex1, although other adaptor roles have previously been suggested. P-Rex1 binding to the actin remodelling protein FLII enhances FLII interaction with Rac1 to control cell/cell contacts, migration and contraction ^49^. As P-Rex1 binds FLII independently of its Rac-GEF activity, this was proposed to be a scaffolding role. However, the Rac-GEF activity of P-Rex1 was required for the downstream effects of FLII, so P-Rex1 still functions as a Rac-GEF in this context. Similarly, P-Rex1 deletion in mice causes a melanoblast migration defect which affects skin pigmentation ^12^. Combined deletion of P-Rex1 and Rac1 brings about a more pronounced phenotype, which suggested that P-Rex1 may have a GEF-independent role in this context ^64^. However, use of GEF-dead P-Rex1 showed that melanoblast migration requires the Rac-GEF activity ^12^, so P-Rex1 likely activates other Rac-type GTPases in addition to Rac1 in this scenario, for example RhoG. It is unsurprising that a protein as large as P-Rex1 should have adaptor functions. Some other Rac-GEFs are also known to play important adaptor roles. For example, Vav family Rac-GEFs have adaptor functions in the NFAT-dependent transcription and integrin-mediated spreading of lymphocytes ^65^, and the Rac-GEF Tiam1 controls dendrite morphology of somatosensory neurons independently of its Rac-GEF activity ^66^.

Our study is not the first to describe a role for P-Rex1 in vesicle trafficking processes. P-Rex1 was shown to be required for the insulin-stimulated upregulation of glucose transporter 4 (GLUT4) from secretory vesicles to the plasma membrane in 3T3-L1 adipocytes, regulating glucose uptake. This could be abolished by the expression of dominant-negative Rac1, suggesting the role of P-Rex1 in GLUT4 trafficking requires its Rac-GEF activity ^67^. Platelets from *Prex1^−/−^* mice have defective dense granule secretion in response to thromboxane stimulation ^68^, and P-Rex1 knockdown in endothelial cells impairs the epinephrine-stimulated secretion of Weibel-Palade bodies ^69^. Further research is needed to elucidate the underlying mechanisms.

We previously identified a direct interaction of P-Rex1 with the GPCR adaptor protein Norbin ^70^, which is mediated through the PH domain and occurs *in vitro* and in cells. Norbin binding increases the catalytic activity of P-Rex1, and both proteins promote each other’s plasma membrane localisation^71^. Norbin binds the C-terminal tails of many GPCRs to regulate GPCR signalling and trafficking ^70^. Norbin also controls S1PR1 trafficking in PC12-S1PR1 cells ^48^, similar to P-Rex1. However, Norbin is not just a positive regulator of P-Rex1, and the roles of P-Rex1 and Norbin in GPCR trafficking appear not to be linked. We showed in mouse neutrophils that Norbin can also function independently of P-Rex1 or even oppose P-Rex1 functions, acting as a suppressor of neutrophil-mediated innate immunity, whereas P-Rex1 is required for this immunity ^8^. We found here that P-Rex1 only affects the agonist-induced internalisation of GPCRs, whereas Norbin largely controls steady-state GPCR trafficking ^70^.

P-Rex1 inhibits the phosphorylation of the C-terminal tails of GPCR which is required for receptor internalisation. P-Rex1 binds Grk2, a major kinase responsible for GPCR phosphorylation, both in cells and *in vitro*, although it does not appear to regulate Grk2 activity. P-Rex1 binding to Grk2 requires the DEP domains, which are central to P-Rex1 regulation and to the downstream transmission of P-Rex1 signals ^3,4^. In addition to Grk2, these domains are also involved in P-Rex1 binding to Gβγ and mTOR, and they harbour S436, a residue phosphorylated by PKA to prevent P-Rex1 activation. However, binding of the DEP domains to Grk2 was not sufficient for the regulation of GPCR trafficking, as the PDZ and IP4P domains of P-Rex1 were also required, whereas the PH domain was dispensable. Others previously suggested that the isolated PDZ domains of P-Rex1 can bind S1PR1 ^52^, but we show that this is not sufficient to affect the trafficking of the receptor. As P-Rex1 does not affect Grk2 activity, we propose that its direct interaction with Grk2 limits the agonist-induced internalisation of GPCRs somehow sterically, for example by preventing access of the Grk2 to the receptor. P-Rex2, the other P-Rex family member, inhibits GPCR trafficking in a similar manner to P-Rex1, and also binds Grk2 in a GEF-activity independent manner. The effect of P-Rex2 on GPCR trafficking appeared less pronounced than with P-Rex1, but this likely reflects lower expression of P-Rex2 in our transient transfections.

The synergistic mode of P-Rex1 activation by PIP_3_ and Gβγ make it an ideal transducer of GPCR signals, and there are numerous examples. Prex1 is required for the fMLP- or C5a-stimulated activation of Rac2, ROS production, actin polymerisation, chemokinesis and chemotaxis in neutrophils ^1,2^, for C5a-and MCP1-stimulated Rac1 activity and chemotaxis in macrophages ^72^, thromboxane A2- and thrombin-dependent secretion of dense granules in platelets ^68^, and SDF1-stimulated Rac1 activity and chemotaxis in endothelial cells ^73^. Furthermore, we previously showed that P-Rex1 is required for the S1P-stimulated Rac1 and Akt activities, spreading and neurite outgrowth in PC12-S1PR1 cells ^47^. The fact that large parts of P-Rex1 are required to control GPCR trafficking hinders the design of mutants which abolish its receptor trafficking role while retaining its signalling capacity. Therefore, it is difficult to dissect how much P-Rex1 mediates GPCR responses through GPCR signalling compared to GPCR trafficking. In light of our findings, we propose that at least in part this will occur through the control of GPCR trafficking, with P-Rex1 preventing the Grk2-depedent desensitisation resulting in retaining high levels of active GPCR at the plasma membrane and therefore prolonging GPCR signalling. Thus, P-Rex1 plays a dual role in promoting GPCR functions.

Finally, our findings have implications for the clinical relevance of P-Rex1 in disease, in particular cancer. P-Rex1 promotes many types of cancer, but like most GEFs, it is difficult to target P-Rex1 directly ^3,4^. In comparison, GPCRs are straightforward targets. Many GPCRs, including S1PR1 and CXCR4, promote cancer growth and metastasis ^74−78^. From our findings, one would predict that cancers with overexpression of P-Rex1 also show high plasma membrane levels of GPCRs, which could be exploited, and therapeutics for these GPCRs may already be approved in the clinic or in development.

## Acknowledgements

We thank Prof. Angeliki Malliri (CRUK Manchester Institute) for the gift of inducible MDCK cells, Prof. Timothy Hla (Harvard University) and Prof. Graham Ladds (University of Cambridge) for GPCR constructs, and Prof. Julie Pitcher (University College London) for Grk2 constructs. We thank the staff of the Babraham imaging facility for their expert help. MB received a PhD studentship from UK Biotechnology and Biological Sciences Research Council (BBSRC). EH received a BBSRC iCASE PhD studentship in collaboration with Vernalis. PI receives a PhD studentship from the Cambridge Trust. EM was awarded a Summer Vacation Studentship from the British Society for Cell Biology. ET received a BBSRC iCASE PhD studentship in collaboration with AstraZeneca. The project was funded by Institute Strategic Programme Grant BB/P013384/1 from the BBSRC to the Babraham Institute Signalling Programme.

## Author contributions

MB, EH, PI, RPM, EM, KH and ET designed, performed and analysed experiments. DH, RH, AM and HW planned and supervised the project and procured funding. MB, EH and HW wrote the manuscript.

## STAR*Methods

### Key resources table

| REAGENT or RESOURCE | SOURCE | IDENTIFIER |
| --- | --- | --- |
| Antibodies |  |  |
| CXCR4 | Novus Biologicals | NB100-56437 |
| EE | Babraham Bioscience Technologies | EE |
| EEA1 | BD Biosciences | 610456 |
| EGFR | Abcam | ab52894 |
| flag | Sigma | F3165 |
| flag agarose-conjugated | Sigma | M8823 |
| GFP | Sigma | G6539 |
| GFP | Abcam | ab290 |
| Grk2 | Cell Signaling Technology | 3982S |
| GST | Merck | Cytiva 27-4577-01 |
| K-Ras | Sigma Aldrich | WH0003845M1 |
| myc | Babraham Bioscience Technologies | myc |
| phospho-S324/S325 CXCR4 | ECM Biosciences | CP435 |
| P-Rex1 | Marcus Thelen <sup>1</sup> | IRB, Bellinzona, Switzerland |
| P-Rex2 | Welch lab <sup>9</sup> | 78 |
| Rab5 | Abcam | ab18211 |
| S1PR1 | Abcam | ab11424 |
| AF568 goat-anti-mouse IgG | Invitrogen | A-11031 |
| HRP donkey anti-goat IgG | Santa Cruz | sc-3851 |
| HRP goat anti-mouse IgG | Bio-Rad | 1706516 |
| HRP goat anti-rabbit IgG | Bio-Rad | 1706515 |
| Bacterial and virus strains |  |  |
| Biological samples |  |  |
| Tubulin purified from pig brain | Tetubio | T240 |
| Chemicals, peptides, and recombinant proteins |  |  |
| Aqua-Poly/Mount | Polysciences, | 18606-20 |
| AY-NH2 | Tocris | 1487 |
| ChromoTek agarose | Proteintech | bmab-20 |
| Dulbecco's Modified Eagle's Medium | Gibco | 41965-039 |
| EGF | Sigma | 11376454001 |
| Escort IV | Sigma-Aldrich | L3287 |
| G418 disulphate | Melford | G0175 |
| GLP-1 | Tocris | 5374 |
| GST | Welch lab <sup>71</sup> | Babraham Institute |
| Hoechst 33342 | Thermo Fisher Scientific | 62249 |
| JetPEI | Polyplus | 101-10N |
| Magnetic high-capacity glutathione agarose | Merck | G0924 |
| OptiPrep solution | StemCell Technologies | 07820 |
| PDGF | Invitrogen | ABC125 |
| Penicillin/streptomycin | Gibco | 15140-122 |
| Protein-A Sepharose | Sigma-Aldrich | P3391 |
| Phosphate Buffered Saline (PBS) | Invitrogen | 70011-036 |
| ProLongGold Antifade | Life Technologies | P36934 |
| Recombinant human wild type and mutant EE-P-Rex1 | Welch lab <sup>1,22,51</sup> | Babraham Institute |
| Recombinant human GST-Grk2 | Abcam | ab125620 |
| S1P | Sigma | S9666 |
| SDF1 | Sigma | SRP3276 |
| Sepharose | Sigma-Aldrich | 4B-200 |
| X-tremeGENE 9 | Roche | 06366511001 |
| Critical commercial assays |  |  |
| ADP-Glo™ kinase assay | Promega | V6930 |
| Clarity western ECL substrate | Bio-Rad | 170-5060 |
| Gateway Bac-to-Bac system | Invitrogen | 11827-011 |
| Site-directed mutagenesis kit | New England Biolabs | E0554 |
| Experimental models: Cell lines |  |  |
| Human Embryonic Kidney 293 (HEK293) cells | Stephens/Hawkins lab | Babraham Institute |
| Madin-Darby Canine Kidney (MDCK) cells with inducible expression of wild type or GEF-dead myc-P-Rex1 | Angeliki Malliri <sup>49</sup> | CRUK Manchester Institute |
| PC12-S1PR1 cells, wild type or knock-out for *Prex1* | Welch lab <sup>47</sup> | Babraham Institute |
| Sf9 insect cells | Stephens/Hawkins lab | Babraham Institute |
| Oligonucleotides |  |  |
| Primers (listed below) | Merck |  |
| Recombinant DNA |  |  |
| pcDNA3-flag-GRK2 | Julie Pitcher | University College London |
| pcDNA3-S1PR1-GFP | Timothy Hla | Harvard University |
| pcDNA3.1-SNAP-GLP1R-mCherry | Graham Ladds | University of Cambridge |
| pcDNA3.1-PAR4-mCherry | Graham Ladds | University of Cambridge |
| pcDNA5FRT-EF-PDGFRβ-eGFP | Addgene | 66790 |
| pCMV3 EE-P-Rex1 and pCMV3 myc-P-Rex1 constructs | Welch lab <sup>1,22,51,79</sup> | Babraham Institute |
| pDEST10 | Invitrogen | 11806-015 |
| pEGFP-N1-EGFR-GFP | Addgene | 32751 |
| pENTR3C | Thermo Fisher Scientific | A10464 |
| pLSSmOrange-N1-hCXCR4-Orange | Addgene | 110197 |
| Software and algorithms |  |  |
| Other |  |  |
| 13 mm glass coverslips | Thermo Scientific | 12392128 |
| 35 mm glass bottom dish | World Precision Instruments | FD35-100 |
| 16 × 102 mm ultra-clear ultracentrifuge tubes | Beckman Instruments | 344661 |
| Immobilon-P PVDF | Millipore | IPVH00010 |
| LoBind tubes | Eppendorf | 0030108132 |
| Nunc T175 Easy flasks | Thermo Fisher Scientific | 159920 |

### Resource availability

#### Lead contact

Further information and requests for resources and reagents should be directed to and will be fulfilled by the lead contact, Heidi Welch.

#### Materials availability

Newly generated materials associated with the paper are available from the lead contact.

#### Data and code availability

- All data reported in this paper will be shared by the lead contact upon request
- This paper does not report original code.
- Any additional information required to reanalyze the data reported in this work paper is available from the lead contact upon request.

### Experimental model and study participant details

Cell lines. Please see Key resources table and Method details sections.

### Method details

#### Expression vectors

Human P-Rex1 cDNA constructs with N-terminal myc or EE epitope tags in pCMV3 were described previously ^1,22,51,79^. mCherry-P-Rex1 was subcloned by replacing the EE-tag of pCMV3(EE)P-Rex1 with mCherry using Kpn1 and EcoR1. pCDNA3-S1PR1-GFP was a gift from Prof. Timothy Hla (Harvard University). pcDNA3.1-SNAP-GLP1R-mCherry and pcDNA3.1-PAR4-mCherry were gifts from Prof. Graham Ladds (University of Cambridge). pLSSmOrange-N1-hCXCR4-Orange (110197), pEGFP-N1-EGFR-GFP (32751) and pcDNA5FRT-EF-PDGFRβ-eGFP (66790) were from Addgene. cDNA3-flag-GRK2 was a gift from Prof. Julie Pitcher (University College London). For the production of recombinant P-Rex2 proteins in Sf9 cells, human P-Rex2 ^31^ was subcloned into pENTR3C. Catalytically inactive (GEF-dead) P-Rex2^E30A,N212A^ was generated in pENTR3C using a site-directed mutagenesis kit (New England Biolabs, E0554) following the manufacturer’s instructions, with primers CGCGTGTGCGTGCTCAGCGCGCTCCAGAAGACCGAGCGG and GCTGTCTGTTCCAACATAGCCGAGGCCAAGAGACAGATG to introduce the E30A and N212A mutations, respectively. The wild type and GEF-dead P-Rex2 clones were recombined with pDEST10 (Invitrogen, 11806-015) to gain an N-terminal 6His tag and generate baculovirus using the Gateway Bac-to-Bac system (Invitrogen, 11827-011).

#### Western blotting

Proteins were transferred onto Immobilon-P PVDF (Millipore, IPVH00010) following SDS-PAGE. Primary antibodies were CXCR4 (Novus Biologicals, NB100-56437, 1:250), EE (clone Glu-Glu, Babraham Bioscience Technologies, 1:50), EEA1 (BD Biosciences 610456, 1:100), EGFR (Abcam, ab52894, 1:1000), flag (clone M2, Sigma, F3165, 1:6000), GFP (Sigma, G6539, 1:2000), Grk2 (Cell Signaling Technology, 3982S, 1:250), GST (Merck, Cytiva 27-4577-01, 1:1000), K-Ras (clone 3B10-2F2, Sigma Aldrich, WH0003845M1, 1:1000), myc (clone 9E10, Babraham Bioscience Technologies, 1:50), phospho-S324/S325 CXCR4 (ECM Biosciences, CP435, 1:250), P-Rex1 ^1^ (clone 6F12, from Prof. Marcus Thelen, IRB, Bellinzona, Switzerland, 1:50), P-Rex2 ^9^ (affinity-purified ‘78’, 1:10000), Rab5 (Abcam, ab18211, 1:1000), and S1PR1 (Abcam, Ab11424, 1:1000). Secondary antibodies were horseradish peroxidase (HRP)-coupled goat anti-rabbit (Bio-Rad, 1706515, 1:3000), goat anti-mouse (Bio-Rad, 1706516, 1:3000) or donkey anti-goat (Santa Cruz, sc-3851, 1:3000). Clarity Western ECL Substrate (Bio-Rad, 170-5060) was used. Where required, membranes were stripped in 25 mM glycine (pH 2.0), 1% SDS for 5 min at RT and reprobed. Coomassie staining (0.1% Coomassie brilliant blue R-250, 50% methanol, 10% acetic acid) of gels and membranes was used to control for protein loading. X-ray films were scanned, and band intensities were quantified by densitometry using Fiji (ImageJ).

#### Cell culture

Mammalian cell lines were used between 1 and 12 weeks in culture. Human Embryonic Kidney 293 (HEK293) cells were grown in Dulbecco’s Modified Eagle’s Medium (DMEM) (Gibco, 41965-039) supplemented with 10% foetal bovine serum (FBS), 100 U/ml penicillin and 100 μg/ml streptomycin (Gibco, 15140-122) at 37°C in a humidified incubator at 5% CO2. To generate HEK293 cells with stable expression of S1PR1-GFP (HEK293-S1PR1 cells), HEK293 cells were transfected with pcDNA.3-S1PR1-GFP using JetPEI, and maintained in the same medium as HEK293 cells except with 500 μg/ml G418 disulphate (Melford, G0175) to select for resistance, and FACS sorted to choose cells with moderate GFP signal. Madin-Darby Canine Kidney (MDCK) cells with doxycline (dox)-inducible expression of myc-tagged wild type or GEF-dead P-Rex1 ^49^ were grown in DMEM with 10% FBS, 100 U/ml penicillin, 100 μg/ml streptomycin, 1 µg/ml puromycin, and 500 µg/ml G418. Expression of wild type or GEF-dead P-Rex1 was induced by adding 1 µg/ml dox for 24 h. PC12 (rat adrenal gland phaeochromocytoma) cells with stable expression of S1PR1-GFP (PC12-S1PR1 cells) which were either wild type or knock-out for *Prex1* ^47^, were grown in poly-D-lysine coated flasks, in DMEM with 10% horse serum, 5% FBS, 100 U/ml penicillin, 100 μg/ml streptomycin, 1× glutamine, and 500 µg/ml G418. Transient transfections were done using JetPEI (Polyplus, 101-10N) or X-tremeGENE 9 (Roche, 06366511001) following the manufacturers’ protocols. Sf9 insect cells for the expression of recombinant proteins were cultured, lipofected using Escort IV transfection reagent (Sigma-Aldrich, L3287), baculovirus particles generated, amplified, and viral titres optimised for protein production as previously described ^51^.

#### GPCR localisation (imaging)

To measure S1PR1 internalisation in HEK293-S1PR1 cells, cells were seeded onto 13 mm coverslips (Thermo Scientific, 12392128) in 24-well plates (Nunc) and transfected the following day with EE-tagged or myc-tagged P-Rex constructs using JetPEI, incubated for 21 h, washed, serum-starved for 6.5 h in DMEM, and then stimulated with 10 nM S1P for various periods of time. Cells were fixed in 4% paraformaldehyde (PFA) in 50 mM Pipes (pH 6.5), 1 mM EGTA, 10 mM MgCl_2_ for 15 min at RT, washed, permeabilised with 0.1% Triton X-100 in PBS for 10 min and washed again. Samples were blocked in PBS/0.5% BSA, incubated with EE antibody (clone Glu-Glu, Babraham Bioscience Technologies, UK, 1:10) or myc antibody (clone 9E10, Babraham Bioscience Technologies, UK, 1:10), washed again, and incubated with goat-anti-mouse AF568-IgG (Invitrogen, A-11031, 1:200), washed in PBS, rinsed in H2O, and mounted using ProLongGold Antifade (Life Technologies, P36934). Cells were imaged using the 60× objective of a Zeiss AxioImager D2 widefield microscope with AxioCam HRm camera. Duplicate coverslips were imaged for each condition and 15 images acquired per coverslip. Images were blinded prior to analysis. To determine the localisation of S1PR1-GFP at the plasma membrane, images were either assessed semi-quantitatively by comparison to a panel of standard images, or were quantified using Volocity or CellProfiler software essentially as described ^48^, by generating a mask covering the entire cell and a second mask shrunk inwards by 0.619 µm (3 pixels), and calculating the GFP signal at the cell edge (mask 1 minus mask 2) as % of the total GFP signal (see also Supplemental Figures 1 and 2).

To measure GPCR internalisation in MDCK cells with inducible expression of P-Rex1 ^49^, cells were seeded onto 13 mm coverslips and transfected the next day using JetPEI to transiently express S1PR1-GFP, GLP1R-mCherry, PAR4-mCherry, or CXCR4-LSSmOrange. Alternatively, EGFR-eGFP or PDGFRβ-eGFP were expressed. The medium was changed 24 h after transfection, and 1 µg/ml dox was added 6 h later to half the samples to induce the expression of wild type or GEF-dead P-Rex1. 24 h after dox treatment, cells were serum-starved for 18 h in DMEM with 100 U/ml penicillin, 100 μg/ml streptomycin, 1 µg/ml puromycin, 500 µg/ml G418, and 0.1% FAF-BSA. The cells were stimulated with the appropriate receptor agonists, namely S1P (Sigma, S9666) for S1PR1, GLP-1 (Tocris, 5374) for GLP1R, AY-NH2 (Tocris, 1487) for PAR4, SDF1 (Sigma, SRP3276) for CXCR4, EGF (Sigma, 11376454001) for EGFR, or PDGF (Invitrogen, ABC125) for PDGFβ, at various concentrations and periods of time, or were mock stimulated. The medium was aspirated, and cells were fixed in 4% PFA for 15 min, washed in PBS, stained with Hoechst 33342 (Thermo Fisher Scientific, 62249, 1:1000), washed again, mounted using ProLongGold Antifade and imaged using the 60× objective of a Nikon AR1 confocal microscope. Receptor localisation was quantified as described here-above.

To measure S1PR1 internalisation in wild type and P-Rex1 deficient PC12-S1PR1 cells, cells were seeded onto 13 mm glass coverslips, serum-starved the next day overnight in DMEM, 0.1% FAF-BSA, and then stimulated with 10 nM S1P for various periods of time, or mock stimulated, and fixed in 4% PFA. Samples washed in PBS, stained with Hoechst 33342, and mounted using Aqua-Poly/Mount (Polysciences, 18606-20). Cells were imaged using the 60× objective of a Nikon AR1 confocal microscope. Duplicate coverslips were assessed per condition, and 5 images acquired per coverslip. Receptor localisation was quantified as described here-above.

#### GPCR localisation (live cell-imaging)

HEK293 S1PR1-GFP cells were seeded into 35 mm glass bottom, dishes (World Precision Instruments, FD35-100) and transfected the following day with mCherry-P-Rex1 using JetPEI. After 21 h, the cells were serum-starved in DMEM for 6 h. Cells were live-imaged using an Olympus CellR widefield imaging system at 37°C, 5% CO2, acquiring frames for GFP and mCherry every 30 s over 45 min. At the flash, an aspirator was used to gently replace the DMEM with DMEM containing 100 nM S1P, keeping a constant volume of 2 ml. Movies were processed using Fiji.

#### GPCR localisation (cell fractionation)

Wild type and P-Rex1 deficient PC12-S1PR1 cells were seeded into poly-D-lysine coated T175 flasks, serum-starved overnight in DMEM, 0.1% FAF-BSA, and then stimulated with 5 nM S1P for 10 min, or mock-stimulated. The medium was aspirated, flasks were transferred onto metal trays on ice, rinsed with ice-cold PBS, and cells harvested by scraping into ice-cold PBS. Cells were centrifuged at 800 × g for 5 min at 4°C, resuspended in 3 ml of ice-cold detergent-free homogenisation buffer (25 mM sucrose, 20 mM Tricine-NaOH, 1 mM EDTA pH 7.8, 2 mM MgCl2, 2 mM DTT, 100 μM PMSF, and 10 μg/ml each of leupeptin, pepstatin-A, aprotinin and antipain) and homogenised in a Teflon-coated homogeniser by douncing. Samples were centrifuged at 800 × g for 10 min at 4°C. 20% OptiPrep solution (StemCell Technologies, 07820) in homogenisation buffer was added to the supernatant to give a concentration of 10% OptiPrep. The sample was loaded onto an OptiPrep step-gradient in 16 × 102 mm ultra-clear ultracentrifuge tubes (Beckman Instruments, Palo Alto, CA, 344661). The step gradient consisted of 5 layers of OptiPrep, 2.3 ml per layer, at 13.3%, 16.6%, 20%, 25% and 40% OptiPrep in homogenisation buffer from top to bottom. Samples were ultracentrifuged for 18 h at 90,000 × g at 4°C in a swinging bucket SW32.1Ti rotor with break/acceleration settings on minimum. 1 ml fractions were collected from each interphase. Proteins were precipitated from the fractions by addition of an equal volume of 25% trichloroacetic acid (TCA) and incubation for 30 min on ice. Samples were centrifuged at 13,225 × g for 15 min at 4°C, and the supernatant was removed. 1 ml of ice-cold acetone was added to each sample, and samples were centrifuged again. The supernatant was removed and the pellet left to air-dry for 20 min. Samples were resuspended in SDS-PAGE sample buffer, with addition of NaOH where necessary to adjust pH, and were analysed by western blotting.

#### Mass spectrometry of S1PR1 phosphorylation

HEK293 S1PR1-GFP cells were seeded into Nunc T175 Easy flasks (Thermo Fisher Scientific, 159920). Half were transfected with pCMV3-EE-P-Rex1 and half with non-expressing control DNA using JetPEI. 21 h later, cells were serum-starved in DMEM for 6.5 h and then stimulated with 10 nM S1P for 10 min, or mock-stimulated. Flasks were transferred onto iced metal trays, washed in PBS, and cells were scraped into 1 ml ice-cold lysis buffer 1 (50 mM Hepes pH 7.2, 150 mM NaCl, 1% Triton X-100, 5 mM EDTA, 0.1 mM PMSF, 1 mM DTT, 20 mM β-glycerol phosphate, 25 mM NaF, 1 mM Na3VO4, 10 μg/ml each of leupeptin, aprotinin, pepstatin-A and antipain). Lysates were centrifuged at 110,000 × g for 30 min at 4°C and the supernatant incubated with 100 μl Sepharose beads (Sigma-Aldrich, 4B-200, prewashed in lysis buffer 1) for 20 min at 4°C with end-over-end rotation. The beads were sedimented at 18,000 × g for 30 s at 4°C and the supernatant transferred into precooled 1.5 ml Eppendorf tubes. 150 μl of supernatant taken as a total lysate sample. 6 μl GFP antibody (Abcam, ab290) was added to the remaining supernatant, and samples were incubated for 1.5 h at 4°C with end-over-end rotation before 60 μl of protein-A Sepharose (Sigma-Aldrich, P3391, prewashed in lysis buffer 1) was added, and samples were incubated for 1 h at 4°C with end-over-end rotation. The beads were sedimented at 18,000 × g for 30 s at 4°C, and the supernatant was removed. 150 μl of the supernatant was kept as a post-immunoprecipitation control. The beads were washed 5 times in lysis buffer 1, protein was eluted by 3 additions of 50 μl 0.1 M glycine, pH 2.5, and the pH was neutralised using 1 M Tris (pH 7.8 at 4°C). The eluates were centrifuged at 18,000 × g for 30 s at 4°C, and the supernatant was transferred to fresh precooled tubes. Boiling 4× SDS-PAGE buffer was added to final 1.3×, and samples were boiled for 10 min and snap-frozen in liquid nitrogen. Samples were subjected to tryptic digest, treated with titanium dioxide to enrich phosphopeptides, and analysed by targeted liquid chromatography mass spectrometry LC-MS. The ratio of phosphorylated to non-phosphorylated peptides was used to quantify the C-terminal phosphorylation of the GPCR.

#### Interaction of P-Rex1 with Grk2 in HEK293-S1PR1 cells

HEK-293-S1PR1-GFP cells were plated into T175 flasks, transfected with myc-P-Rex1 and/or flag-Grk2 using jetPEI for 72 h, and then serum-starved in DMEM for 14 h. Cells were washed in PBS (Invitrogen, 70011-036), scraped, centrifuged at 10,000 × g for 30 s at 4°C, and resuspended in ice-cold lysis buffer 2 (50 mM Hepes, pH 7.2 at 4°C, 150 mM NaCl, 1% NP-40, 1 mM EDTA, 2 mM EGTA, 1 mM DTT, 0.1 mM PMSF, and 25 μg/ml each of leupeptin, pepstatin-A, aprotinin, and antipain). The lysate was incubated on ice for 10 min with intermittent vortexing, cleared by centrifugation at 10,000 × g for 3 min at 4°C, and the supernatant recovered. For a total lysate control, boiling 4× SDS-sample buffer was added to 75 μl of cleared lysate, samples boiled for 5 min, and frozen in liquid nitrogen. The rest of the lysate was transferred into precleared 2 ml LoBind tubes (Eppendorf, 0030108132) and precleared with 10 µg prewashed magnetic ChromoTek agarose (Proteintech, bmab-20) for 30 min at 4°C with end-over-end rotation. The beads were removed magnetically, and the supernatant was transferred into fresh tubes and incubated with 10 µg prewashed magnetic anti-flag agarose beads (Sigma, M8823) for 60 min at 4°C with end-over-end rotation. 75 µl of the supernatant was retained as a ‘supernatant’ control and processed like the total lysate control. The beads were washed 4 times and resuspended in 30 µl boiling 1.3× SDS-sample buffer, boiled for 5 min, and frozen in liquid nitrogen. Samples were analysed by western blotting with P-Rex1 and flag antibodies.

Co-immunoprecipitation of P-Rex1 with S1PR1-GFP was performed the same way, except that HEK293-S1PR1 cells were transfected with myc-P-Rex1 alone, or mock-transfected, and stimulated with 100 nM S1P for 10 min, or mock-stimulated, after the serum-starvation, S1PR1-GFP was immunoprecipitated using magnetic GFP-trap agarose, and samples were analysed by western blotting with myc and S1PR1 antibodies.

#### Recombinant proteins

Recombinant human wild type and mutant EE-P-Rex1 proteins, purified from baculovirus-infected Sf9 cells using their EE tag, were as previously described ^1,22,51^. Recombinant human wild type and GEF-dead His-P-Rex2 proteins were purified from baculovirus-infected Sf9 cells using their His tag. Pellets from 400 ml Sf9 cell cultures infected with high titre baculovirus (see above) were thawed into 25 ml ice-cold lysis buffer 3 (PBS, 1% Triton X-100, 25 mM NaF, 20 mM β-glycerophosphate, 1 mM DTT, 0.1 mM PMSF and 10 μg/ml each of antipain, pepstatin A, leupeptin, aprotinin), lysed on ice for 5 min, and ultracentrifuged at 200,000 × g for 1 h at 4°C. The supernatant was incubated with prewashed Ni-NTA agarose for 90 min at 4°C and with end-over-end rotation. Beads were washed 3 times in ice-cold 2× PBS, 1% Triton X-100 and 4 times in wash/elution buffer (PBS, 10% glycerol, 1 mM DTT, 0.01% azide, 20 mM imidazole). For elution, wash/elution buffer containing 600 mM imidazole was added, and samples were incubated for 10 min on ice. Samples were centrifuged at 800 × g for 1 min at 4°C and the supernatant recovered. A second elution was performed with 300 mM imidazole and pooled with the first. To remove the imidazole, a PD-10 desalting column (GE Healthcare Life Sciences, 52130800) was used according to the manufacturer’s instructions. Desalted protein was concentrated using a 100 kDa Amicon Ultra filter (Merck, UCF210024). Once the sample volume was reduced to 100 μl, 2 ml of equilibration buffer (1× PBS, 10% glycerol, 1 mM DTT, 1 mM EGTA, 0.01% azide) were added, and the samples concentrated again. Glycerol was added to 50%, and protein for GEF activity assays was supplemented with 2 mg/ml FAF-BSA. To test the quality of the purified P-Rex2 proteins, Rac-GEF activity was measured using a liposome-based assay as previously described ^1,31,5^. GST was purified from *E. coli* as previously described ^71^. Sf9 cell-derived recombinant human GST-Grk2 was from Abcam (Abcam, ab125620).

#### Direct binding of P-Rex proteins to Grk2

5 pmol of P-Rex1 or P-Rex2 protein were incubated with 5 pmol of either GST or GST-Grk2 in a volume of 20 µl in detergent-free buffer (50 mM Hepes, pH 7.2 at 4°C, 150 mM NaCl, 1 mM EDTA, 2 mM EGTA, 1 mM DTT, 0.1 mM PMSF, and 25 μg/ml each of leupeptin, pepstatin-A, aprotinin, and antipain) for 1 h on ice, with frequent vortexing. For experiments with P-Rex1 mutants, proteins were reduced to 2.5 pmol. A 2 µl aliquot was taken as the ‘reaction mix’ control. Boiling 1.3x SDS-sample buffer was added, and the sample was boiled for 5 min, and frozen in liquid nitrogen. The remaining reaction mix was added to 300 μl detergent-free buffer in LoBind tubes containing 5 μl prewashed magnetic high-capacity glutathione agarose (Merck, G0924) and incubated for 45 min at 4°C with end-over-end rotation. The beads were sedimented using a magnet, and 75 μl of the supernatant was retained as a ‘supernatant’ control, which was processed like the ‘reaction mix’ control. The beads were washed four times in lysis buffer 2, boiling 1.3× SDS-sample buffer was added, and samples were boiled for 5 min, and frozen in liquid nitrogen. Samples were analysed by western blotting using P-Rex1 or P-Rex2 and GST antibodies.

#### Grk2 kinase activity

To measure the catalytic activity of Grk2, the ADP-Glo™ kinase assay kit (Promega, V6930) was used with tubulin as the substrate. 40 nM human recombinant GST-Grk2 and/or 40 nM EE-P-Rex1 proteins were incubated with 150 nM tubulin purified from pig brain (Tetubio, T240) and 400 µM ATP in kinase buffer (40 mM Tris, pH 7.5 (RT), 20 mM MgCl_2_, and 0.1% BSA), in a volume of 25 µl for 30 min at 30°C. Controls included samples without protein and with kinase detection reagent only. To control for potential effects of the storage buffers of EE-P-Rex1 (PBS, 1 mM EGTA, 1 mM DTT, 50% glycerol, 0.01% sodium azide) and GST-Grk2 (0.79% Tris HCl, 0.88% NaCl, 0.31% glutathione, 0.002% PMSF, 0.004% DTT, 0.003% EDTA, 25% glycerol), the buffers were added to samples without protein at the equivalent dilution. After the incubation, 25 µl ADP-Glo reagent was added for 40 min at RT to deplete any remaining ATP. 50 µl of kinase detection reagent was added for a further 40 min, and luminescence was measured in a PHERAstar FS luminometer (BMG Labtech).

### Quantification and statistical analysis

Data were tested for normality of distribution to determine if parametric or non-parametric methods of analysis were appropriate. For comparison of two groups, unpaired Student’s t-test was used, whereas for comparison of multiple groups, one-way or two-way ANOVA was used, as appropriate, with repeated measures followed by post-hoc test with multiple comparisons correction. Parameters with values of p ≤ 0.05 were considered to differ significantly. In the figures, * indicates p < 0.05, ** p < 0.01, *** p < 0.001, and **** p < 0.0001. Results are presented as mean ± standard error of the mean (SEM). The number of experimental repeats is indicated in the figure legends. Statistical analysis and plotting of graphs were performed in GraphPad Prism 10.

