## Supplemental figures 1-7 and legends for "P-Rex1 Limits the Agonist-Induced Internalisation of GPCRs Independently of its Rac-GEF Activity"

[4] Bioscience Metabolism, Research and Early Development, Cardiovascular, Renal and Metabolism (CVRM), BioPharmaceuticals R&D, AstraZeneca, Cambridge, UK

[5] Vernalis (R&D) Ltd., Granta Park, Cambridge, UK

**Contents:**

Supplemental Figures 1-7

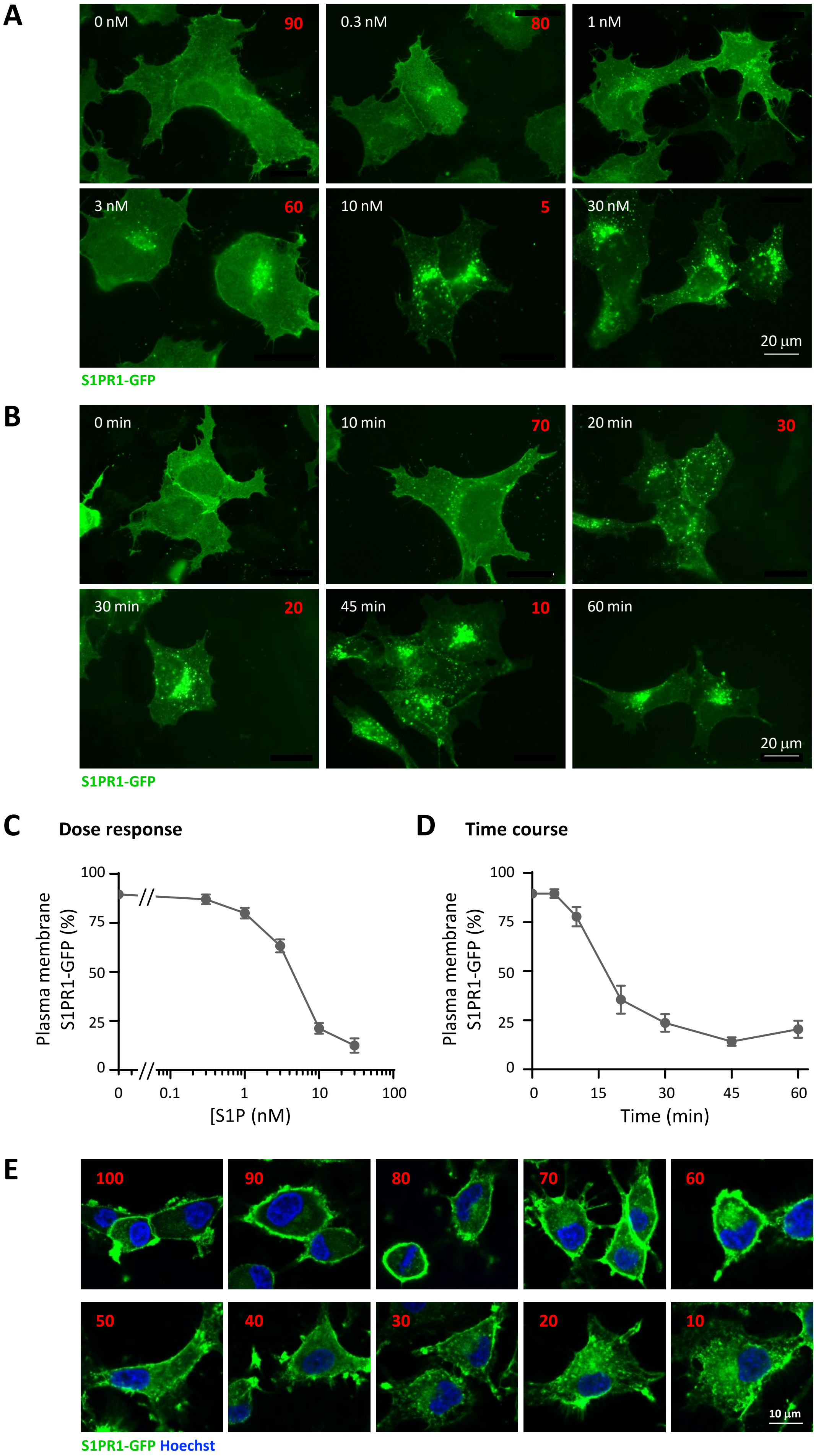


Supplemental Figure 1

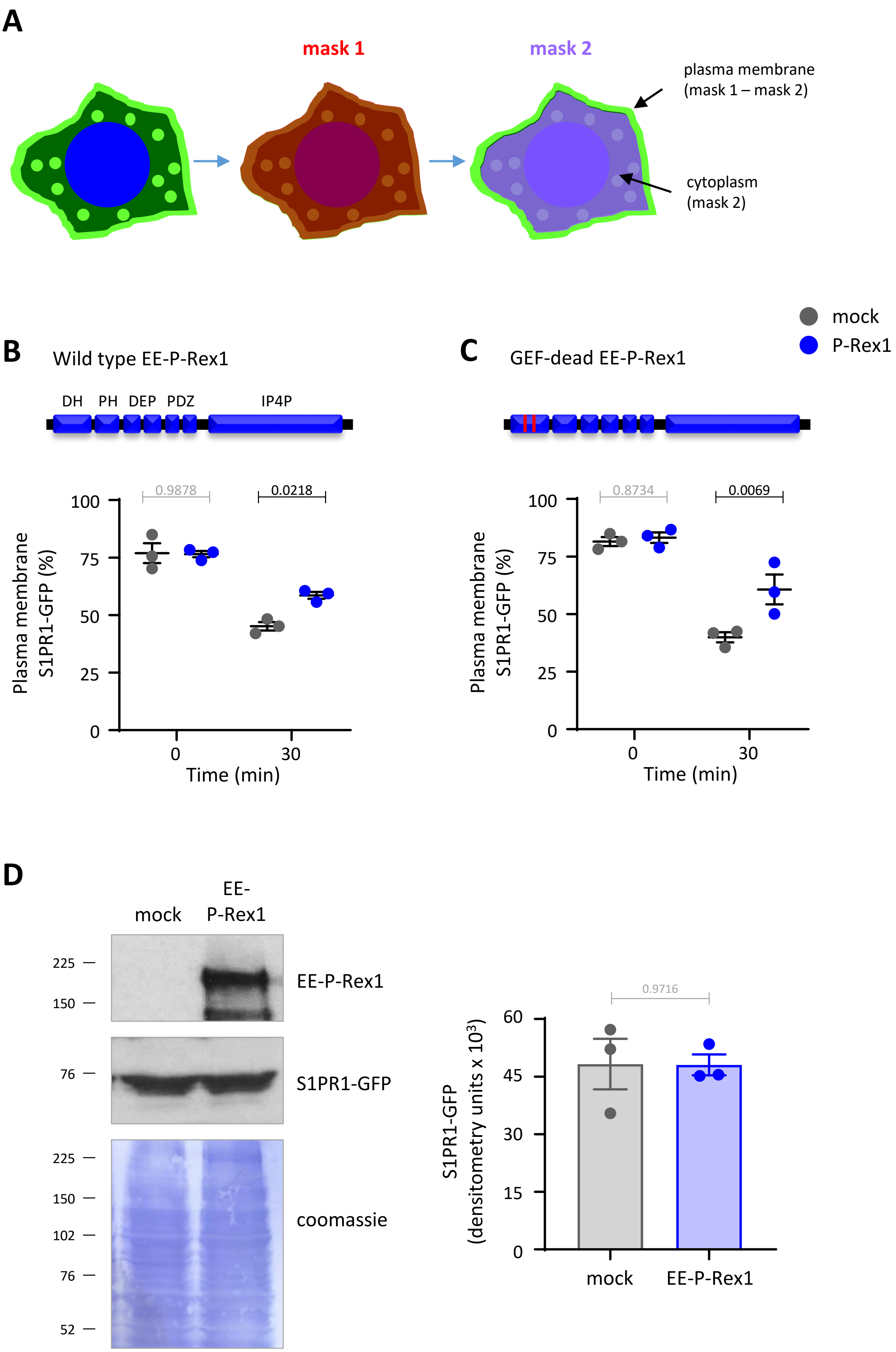


Supplemental Figure 2

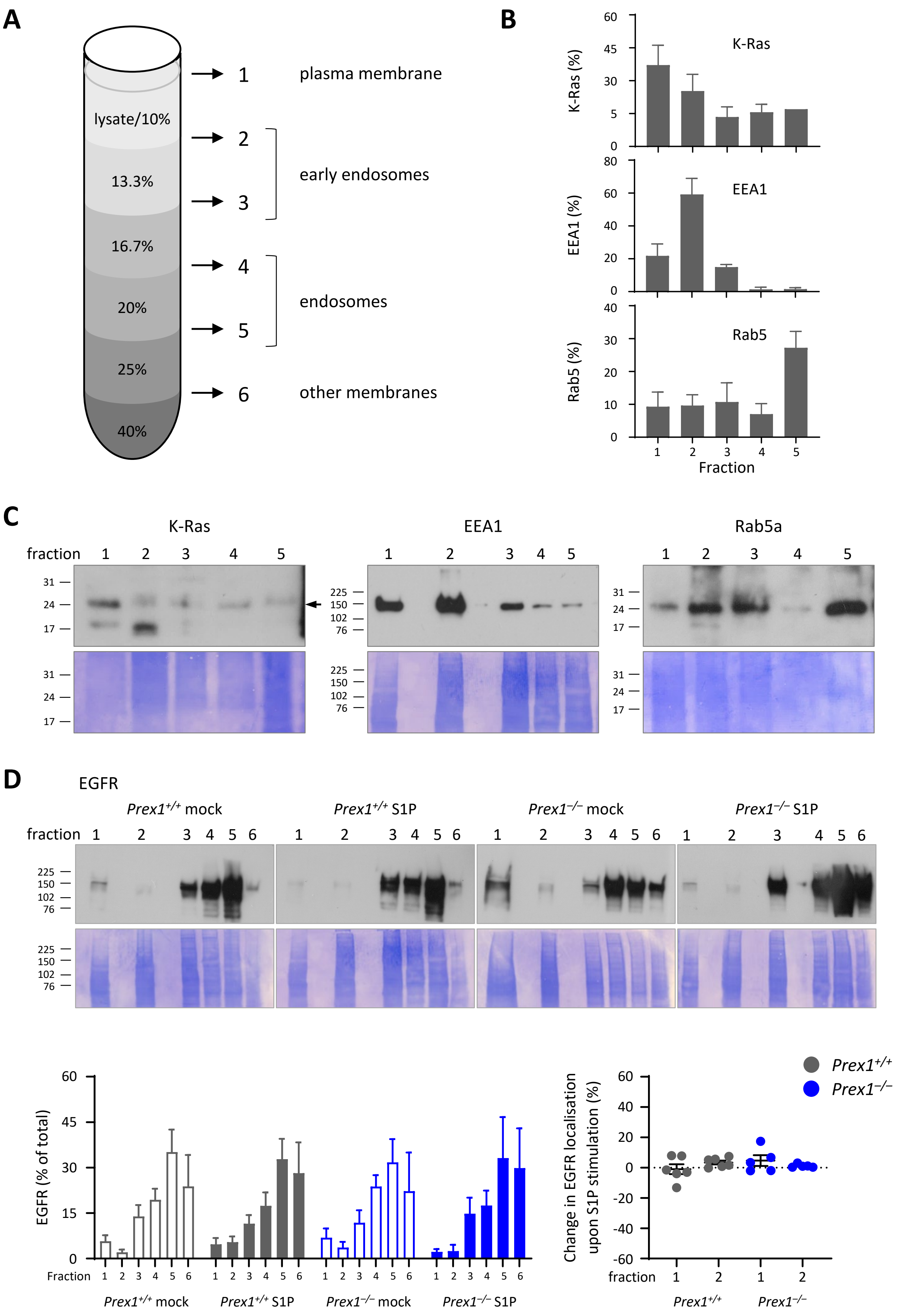


Supplemental Figure 3

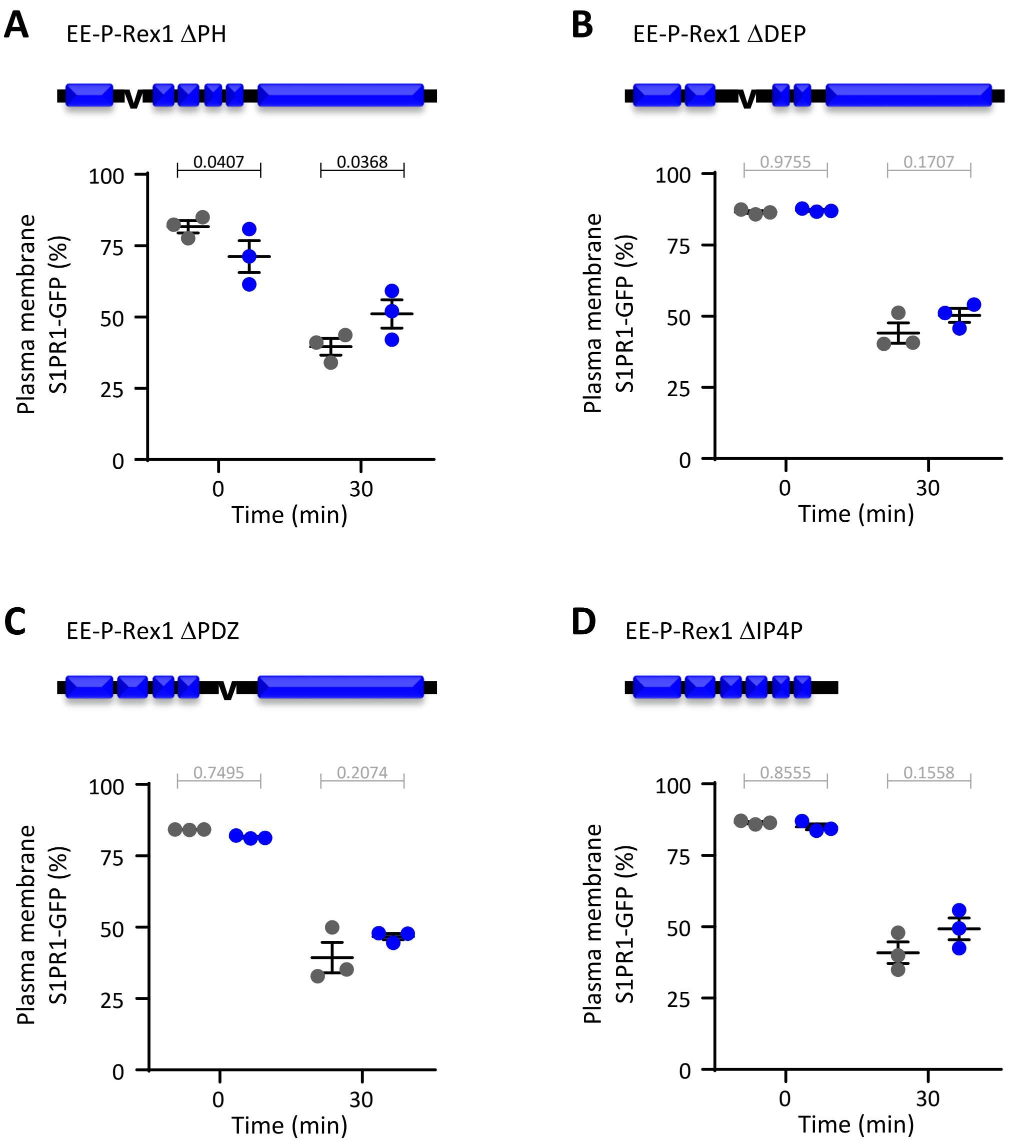


Supplemental Figure 4

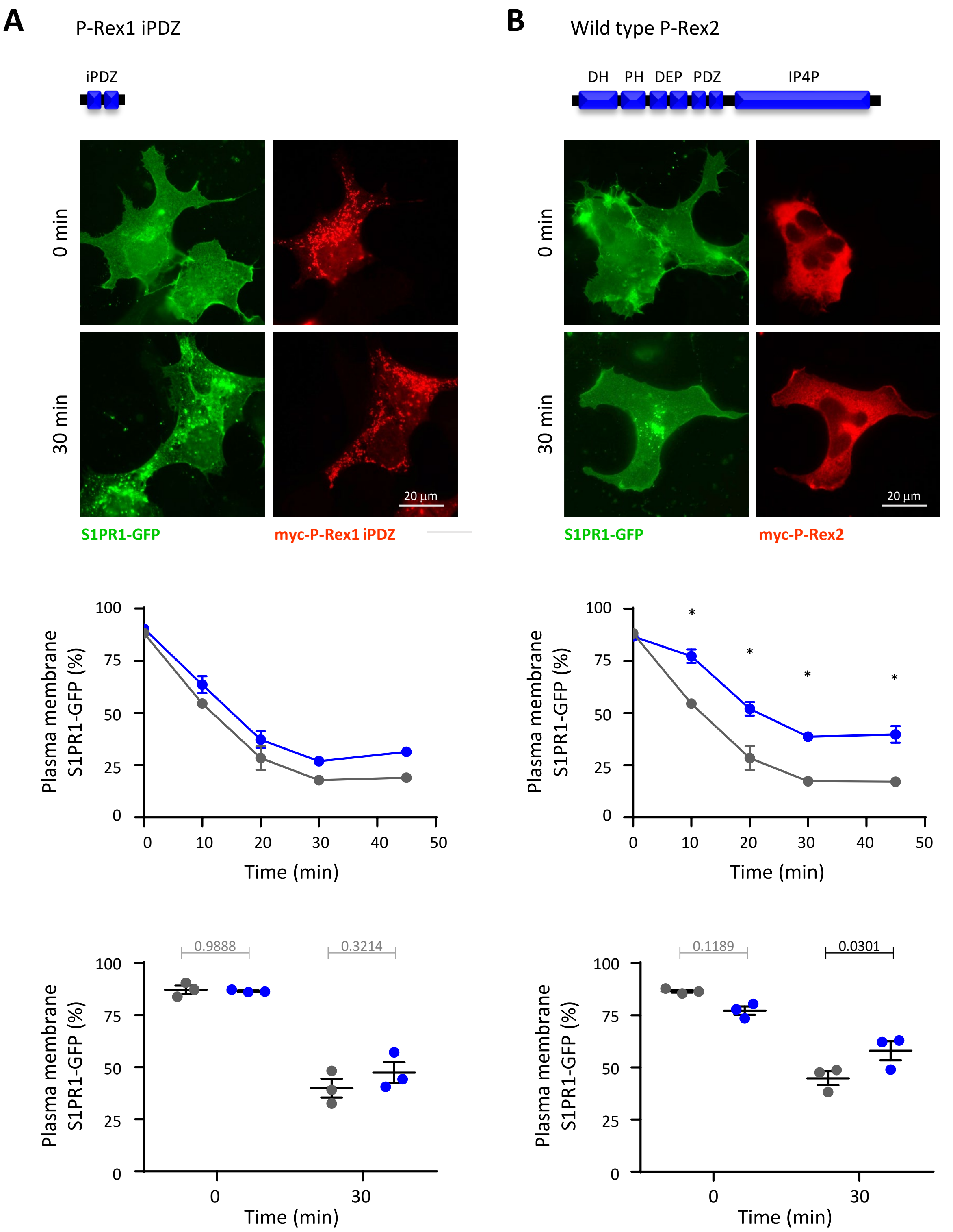


Supplemental Figure 5

A

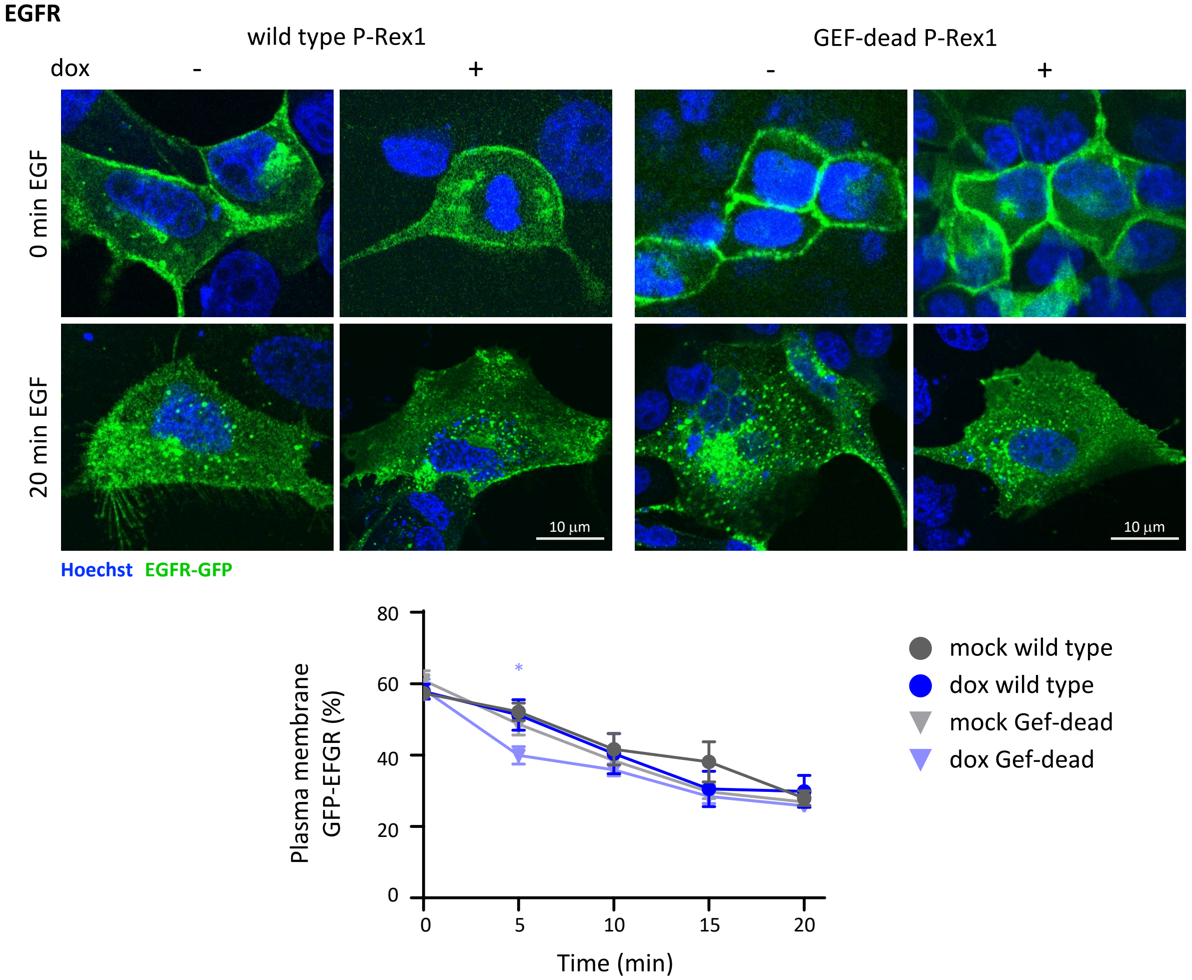


B

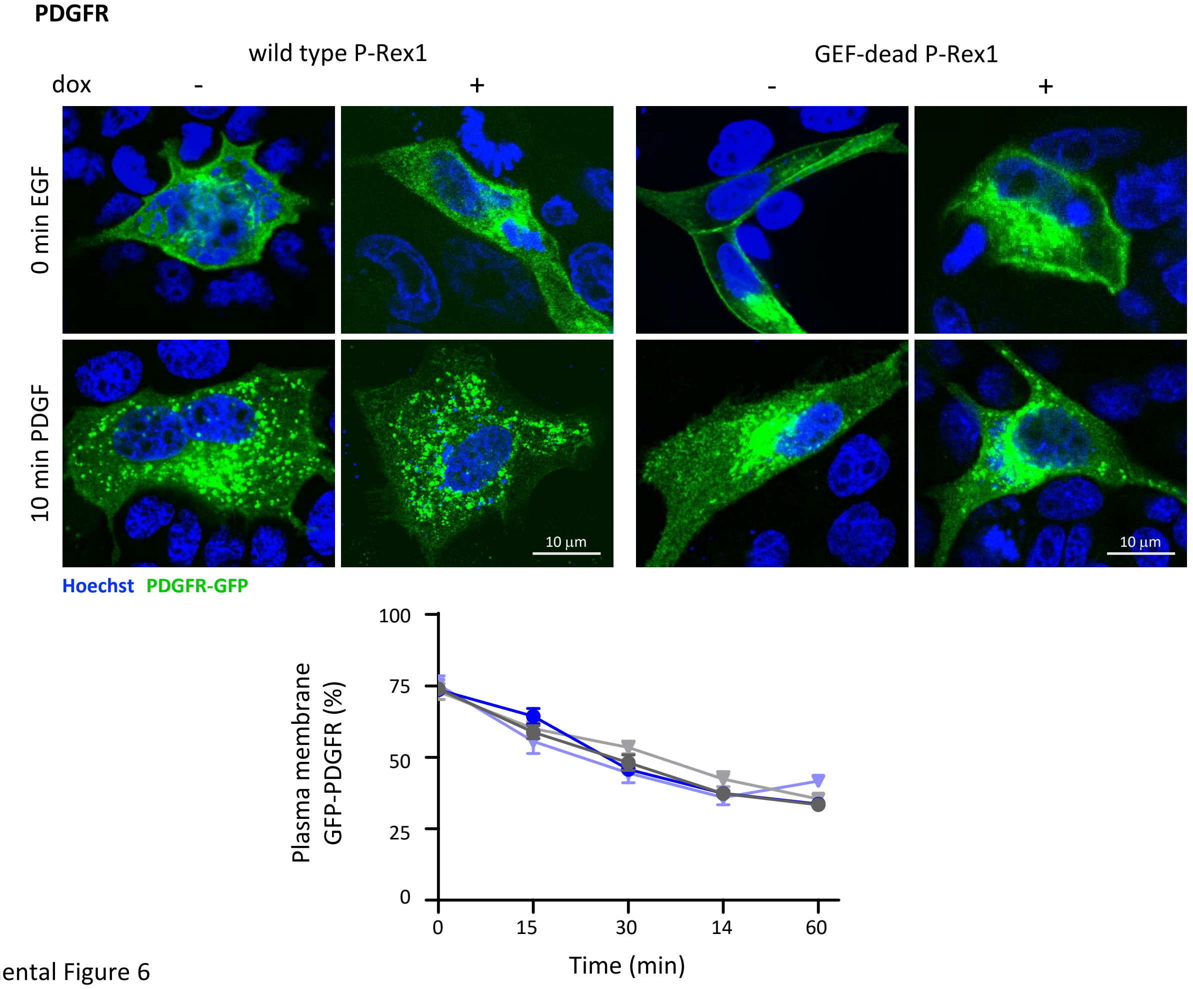


Supplemental Figure 6

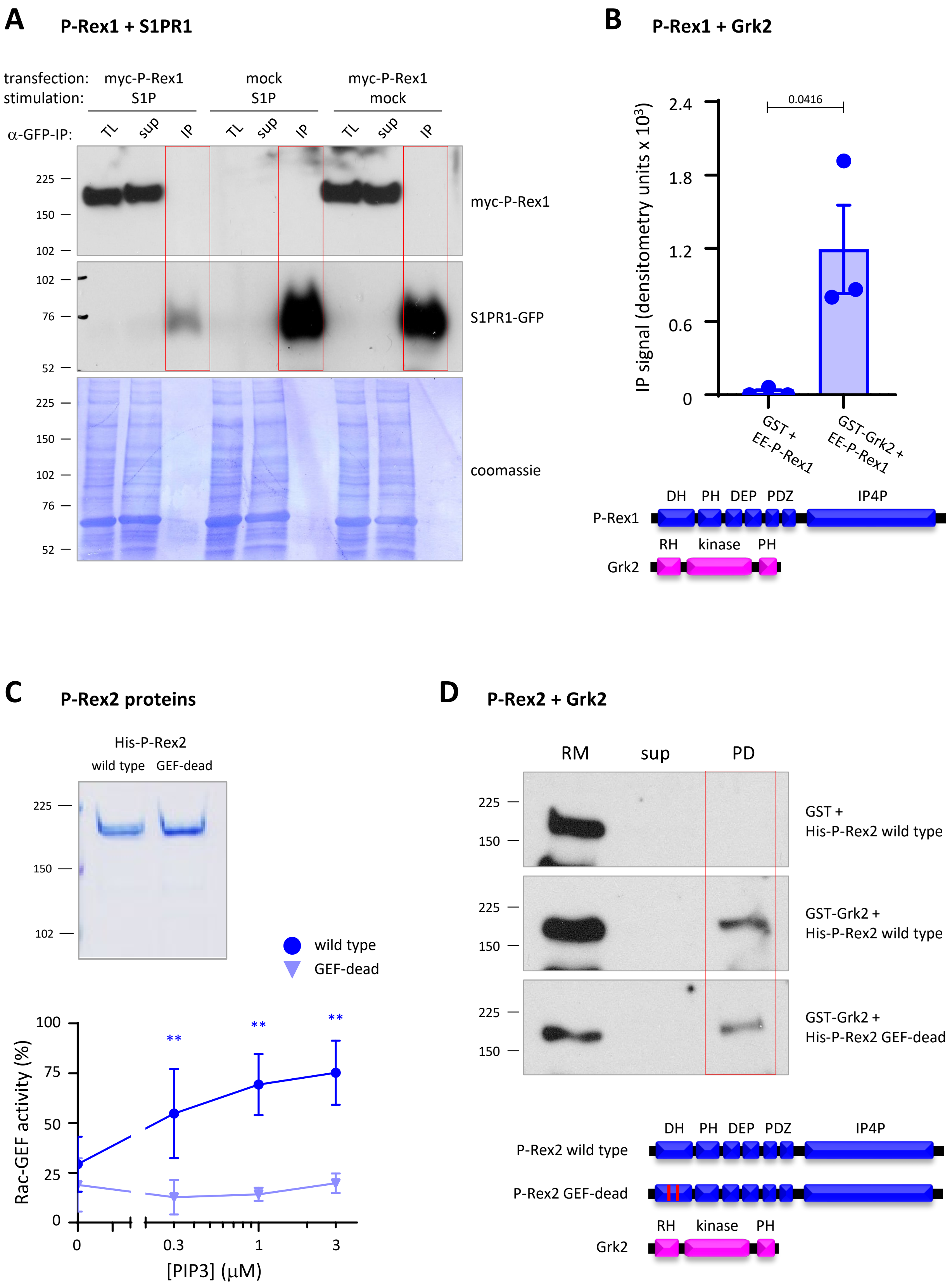


Supplemental Figure 7

**Supplemental Figure Legends**

**Supplemental Figure 1. S1PR1-GFP is internalised in a dose- and time-dependent manner upon S1P stimulation of HEK293-S1PR1 and PC12-S1PR1 cells.** HEK293-S1PR1 cells were serum-starved and stimulated with the indicated concentrations of S1P for 30 min **(A, C)** or stimulated with 10 nM S1P for the indicated periods of time **(B, D)**, fixed and imaged by widefield fluorescence microscopy. (A,B) show representative images, which show the sheet-like appearance of S1PR1-GFP typical for plasma membrane proteins imaged by widefield microscopy and the vesicular localisation in S1P-stimulated cells. Images with red numbers (in %) form part of a panel of standard images used to determine how much receptor is localised at the plasma membrane. (C,D) Quantification of S1PR1-GFP at the plasma membrane by comparison of blinded images to a panel of standard images. Data are mean ± SEM of cells from one of two similar pilot experiments. **(E)** Representative images for comparison of S1PR1-GFP localisation in PC12-S1PR1 cells. PC12-S1PR1 were serum-starved and stimulated for 10 min with S1P concentrations ranging from 0-50 nM, fixed, stained with Hoechst 33342, and imaged by confocal fluorescence microscopy. Images show the S1PR1-GFP ring at the cell periphery typical for plasma membrane proteins imaged by confocal microscopy, and the vesicular localisation in S1P-stimulated cells. Red numbers (in %) denote the amount of S1PR1-GFP at the plasma membrane.

**Supplemental Figure 2. P-Rex1 limits the S1P-dependent internalisation of S1PR1 independently of its catalytic Rac-GEF activity. (A)** Schematic showing quantification of receptor localisation at the plasma membrane by Volocity or CellProfiler image analysis. Mask 1 is generated to cover the entire cell and shrunk inwards by 0.619 µm (3 pixels) to generate mask 2. The fluorescent signal at the cell edge (mask 1 minus mask 2) is calculated as % of the total. **(B, C)** Expression of wild type or GEF-dead P-Rex1 limits the S1P-induced internalisation of S1PR1 in HEK293-S1PR1 cells. HEK293-S1PR1 cells were transfected with wild type (B) or GEF-dead (C) EE-P-Rex1 (blue symbols), or mock transfected (grey symbols), serum-starved, and stimulated with 10 nM S1P for 0 or 30 min, fixed and stained with EE antibody. The amount of S1PR1-GFP at the plasma membrane was quantified by Volocity image analysis as in (A). Data are mean ± SEM of 3 independent experiments, the same as those shown in Figures 1A and 1B. Statistics are two-way ANOVA with Sidak's multiple comparisons correction; p-values in black are significant, p-values in grey are not. **(D)** P-Rex1 does not affect total S1PR1-GFP levels. Total lysates of HEK293-S1PR1 cells expressing EE-P-Rex1, or mock-transfected, were western blotted with P-Rex1 and GFP antibodies. Coomassie staining was used to control for protein loading. Western blots were quantified by Fiji densitometry. Data are mean ± SEM of 3 independent experiments. Statistics are paired t-test; p-values in grey are not significant.

**Supplemental Figure 3. P-Rex1 deficiency does not affect EFGR localisation in S1PR1-GFP cells. (A-C)** Validation of cell fractionation method. PC12-S1PR1 cells were serum-starved, stimulated with 5 nM expressed. Quantification of S1PR1-GFP localisation was done as in (A). Data are mean ± SEM of 3 independent experiments. Statistics are two-way ANOVA with Sidak's multiple comparisons correction; stars denote differences between conditions with and without P-Rex2 for each time point.

**Supplemental Figure 6. P-Rex1 does not control the agonist-induced internalisation of the RTKs EGFR and PDGFR. (A)** EGFR. EGFR-GFP was expressed in MDCK cells with dox-inducible expression of wild type (circles) or GEF-dead (triangles) P-Rex1. Cells were treated with 1 µg/ml dox (blue symbols) for 24 h, or mock-treated (grey symbols), serum starved, and then stimulated with 100 ng/ml for the indicated periods of time, fixed, stained with Hoechst 33342, and imaged by confocal fluorescence imaging. Representative confocal images are shown. EGFR-GFP localisation was quantified by comparison to standard images (see Supplemental Figure 1). **(B)** PDGFR. MDCK cells were treated as in (A) except that PDGFRβ-GFP was expressed and cells were stimulated with 40 ng/ml PDGF. PDGFRβ-GFP localisation was quantified as in (A). Data in (A, B) are mean ± SEM of three independent experiments for each receptor. Statistics are two-way ANOVA with Sidak's multiple comparisons correction.

**Supplemental Figure 7. P-Rex1 does not interact with S1PR1-GFP, but binds GRK2, as does P-Rex2. (A)** HEK293-S1PR1 cells were transfected with myc-P-Rex1, or mock-transfected, serum-starved, stimulated with 100 nM S1P, or mock-stimulated, for 10 min, lysed, subjected to immunoprecipitation (IP) with GFP antibody, and analysed by western blotting with myc and S1PR1 antibodies. 1.5% of the total lysate (TL) and IP supernatant (sup) and all of the IP sample were loaded. Coomassie staining was used as a loading control. Blots shown are representative of three independent experiments. **(B)** P-Rex1 binds directly to Grk2 *in vitro*. Myc-P-Rex1 was incubated with GST or GST-Grk2, proteins were isolated using GSH-beads, western blotted with P-Rex1 and GST antibodies, and quantified using Fiji densitometry. Representative blots are shown in Figure 6B. Data are mean ± SEM of three independent experiments; statistics are paired t-test. **(C)** Human recombinant wild type His-P-Rex2 (blue bars) and mutated His-P-Rex2$^{E30A,N212A}$ (GEF-dead) were produced in Sf9 cells and purified using Ni-NTA agarose. A coomassie-stained gel of the purified proteins is shown. To confirm that wild type His-P-Rex2 is active and GEF-dead His-P-Rex2 is not, the purified proteins were tested in a liposome-based *in vitro* GEF activity assay, using GDP-loaded EE-Rac1 as the substrate and the indicated doses of $PIP_3$ to activate P-Rex2. Rac-GEF activity ($^{35}$S-GTPγS loading of EE-Rac1) is quantified as % of a positive control containing EDTA. Data are mean ± SEM of 3 independent experiments; statistics are two-way ANOVA with Sidak's multiple comparisons correction; stars denote differences for each $PIP_3$ concentration. **(D)** P-Rex2 binds directly to Grk2 *in vitro*, independently of its Rac-GEF catalytic activity. Wild type or GEF-dead P-Rex2 proteins as in (C) were incubated with GST or GST-Grk2, isolated using GSH-beads, and western blotted with P-Rex2 and GST antibodies. 10% of the reaction mix (RM) and pull-down supernatant (sup) controls and all of the pull down (PD) sample were loaded. Blots are representative of 3 independent experiments.

**Supplemental Movie Legends**

**Supplemental Movie 1. S1P-stimulated internalisation of S1PR1-GFP in HEK293-S1PR1 cells.** HEK293 S1PR1-GFP cells were serum-starved and live-imaged in an Olympus CellR widefield imaging system for 45 min, acquiring frames for GFP every 30 s. At the flash, 100 nM S1P was added. The representative movie is from one of four independent experiments.

**Supplemental Movie 2. Expression of mCherry-P-Rex1 inhibits the S1P-stimulated internalisation of S1PR1-GFP in HEK293-S1PR1 cells.** HEK293 S1PR1-GFP cells were transiently transfected to express mCherry-P-Rex1, serum-starved and live-imaged in an Olympus CellR widefield imaging system for 45 min, acquiring frames for GFP and mCherry every 30 s. At the flash, 100 nM S1P was added. The representative movie is from one of six independent experiments.
